# Localization of the Lens Intermediate Filament Switch by Imaging Mass Spectrometry

**DOI:** 10.1101/2020.04.21.053793

**Authors:** Zhen Wang, Daniel J. Ryan, Kevin L Schey

## Abstract

Imaging mass spectrometry (IMS) enables targeted and untargeted visualization of the spatial localization of molecules in tissues with great specificity. The lens is a unique tissue that contains fiber cells corresponding to various stages of differentiation that are packed in a highly spatial order. The application of IMS to lens tissue localizes molecular features that are spatially related to the fiber cell organization. Such spatially resolved molecular information assists our understanding of lens structure and physiology; however, protein IMS studies are typically limited to abundant, soluble, low molecular weight proteins. In this study, a method was developed for imaging low solubility cytoskeletal proteins in the lens; a tissue that is filled with high concentrations of soluble crystallins. Optimized tissue washes combined with on-tissue enzymatic digestion allowed successful imaging of peptides corresponding to known lens cytoskeletal proteins. The resulting peptide signals facilitated segmentation of the bovine lens into molecularly distinct regions. A sharp intermediate filament transition from vimentin to lens-specific beaded filament proteins was detected in the lens cortex. MALDI IMS also revealed the region where posttranslational myristoylation of filensin occurs and the results indicate that truncation and myristoylation of filensin starts soon after filensin expression increased in the inner cortex. From intermediate filament switch to filensin truncation and myristoylation, multiple remarkable changes occur in the narrow region of lens cortex. MALDI images delineated the boundaries of distinct lens regions that will guide further proteomic and interactomic studies.

## 1. Introduction

The ocular lens is a unique tissue that requires precise cell-cell organization to perform its normal function as a transparent, refractory optical element. From a monolayer of lens epithelial cells to fiber cells in distinct stages of differentiation, high spatial order of fiber cell organization is maintained. High resolution microscopy together with immunohistochemistry has allowed significant progress to be made in our understanding of fiber cell organization (Costello et al., 2013; Farnsworth et al., 1974; Lim et al., 2009; Taylor et al., 1996; Vrensen et al., 1992). Revealed by electron microscopy, fiber cells in different lens regions, from outer cortex to inner nucleus, have characteristic cell shapes and size (Farnsworth et al., 1974; Taylor et al., 1996). Ultrastructure studies also revealed high spatial organization of complex membrane architecture and interdigitations along with the maturation and compaction of lens fiber cells (Audette et al., 2017; Kuwabara, 1975; Lo and Harding, 1984; Lo et al., 2014; Taylor et al., 1996; Vrensen et al., 1992). Recently, with better preservation of the epithelium and outer cortex of the lens, two unique zones of membrane remodeling (remodeling zone and transition zone) within the outer cortex region were characterized in both human and nonhuman primate lenses (Costello et al., 2013, 2014; Lim et al., 2009). This structural transformation is expected to be accompanied by changes in proteins and lipids (Vrensen et al., 1992). Knowledge of the spatial distributions of different biological molecules is important for understanding the regulation of cell organization in the lens that will help to develop treatments for lens diseases that are caused by disruption of cellular packing and intracellular connections.

Imaging mass spectrometry (IMS) is a technology that enables the untargeted analysis of thousands of molecules from the surface of a tissue section without the need for labels or antibodies (Caprioli et al., 1997; Stoeckli et al., 2001). IMS has been utilized to characterize a number of tissue types (Buchberger et al., 2018) and has been applied to study the distribution of lens crystallins and their phosphorylation and truncation products in lens tissue (Grey and Schey, 2009a, 2008; Stella et al., 2010). Optimization of washing and instrument conditions allowed imaging of the lens major membrane protein AQP0 (Grey et al., 2009b) and lens β- and γ-crystallins (Anderson et al., 2019). Posttranslational modifications such as deamidation, truncation, and fatty acylation of AQP0 in different lens regions and in different aged lenses have also been investigated (Schey et al., 2010; Wenke et al., 2015). IMS was also applied to charaacterize lens lipids and metabolites (Deeley et al., 2010; Nye-Wood et al., 2017; Seng et al., 2018; Vidová et al., 2010).

Although MALDI IMS is a rapidly developing field with dramatic improvements in instrument mass resolving power and scan speed introduced (Prentice et al., 2018; Spraggins et al., 2016), and the improvement of spatial resolution (Spivey et al., 2019), there are particular challenges for protein imaging (Heeren et al., 2009). Specifically, imaging and identification of low abundance and high molecular weight proteins is challenging (Ryan et al., 2019a). The ocular lens is filled with high concentrations of crystallins (up to 450 mg/mL) (Vérétout et al., 1989) and these high concentrations can suppress the ionization of lower abundance proteins, limiting their detection from tissue. The sensitivity of IMS for intact proteins is generally low and intact protein imaging is limited to abundant proteins and small proteins with a molecular weight less than ∼30 kDa (Franck et al., 2010). In addition, soluble proteins are more favored in the MALDI process (Seeley and Caprioli, 2008); however, many important proteins are neither small nor soluble. To overcome these limitations, *in situ* digestion protocols were developed that typically yield peptides between 400 and 3500 Da; a range in which instrumental sensitivity is higher than that for intact proteins (Groseclose et al., 2007; Judd et al., 2019; Lemaire et al., 2007).

Other than highly abundant crystallins, the lens has many other important proteins that have molecular weights greater than 30 kDa including many cytoskeletal proteins and lens membrane proteins. Three lens intermediate filament proteins, beaded filament structural protein 1 (BFSP1 or filensin), beaded filament structural protein 2 (BFSP2, CP49 or phakinin), and vimentin, are highly abundant and have been shown to be essential for lens optical properties and maintaining lens transparency (FitzGerald, 2009; Song et al., 2009). Vimentin is expressed in lens epithelium and young fiber cells, and the expression of vimentin is dramatically reduced in older fiber cells (Sandilands et al., 1995b) and completely disappears in the center of the lens (Ellis et al., 1984; Sandilands et al., 1995b). CP49 and filensin are believed to form a distinct network, independent of the vimentin network, and changes in subcellular distribution were found to be correlated with organelle loss (Sandilands et al., 1995b). However, when studying the superficial fiber cells, an intermediate filament switch from vimentin to lens beaded filaments was found to be associated with the formation of a remodeling zone in human lenses (Wenke et al., 2016).

Herein, we describe a new washing method that was optimized for retaining intermediate filament proteins, one of the most insoluble cytoskeletal structures of eukaryotic cells (Nicholl and Quinlan, 1994), within lens tissue, while removing the abundant crystallin proteins. MALDI imaging of these proteins at the peptide level after *in situ* digestion allowed visualization of their spatial distribution across the lens. Our results suggest that multiple intermediate filament remodeling processes may occur in the superficial cortex about 1 mm interior to the lens from the capsule.

## 2. Materials and Methods

### 2.1. Materials

Trifluoroacetic acid (TFA), urea, Triton X-100, α-cyano-4-hydroxycinnamic acid (CHCA) and 2, 5-dihydroxybenzoic acid (DHB) were purchased from Sigma-Aldrich Corp. (St. Louis, MO). CHCA and DHB were recrystallized before use. Bovine lenses were purchased from Pel-Freez Biologicals (Rogers, AR) and stored at −80°C until further use. MS grade trypsin, GluC and Lys C protease were purchased from Thermo Scientific (Rockford, IL). Ammonium formate, formic acid, high-performance liquid chromatography (HPLC)-grade acetonitrile and water were purchased from Fisher Scientific (Pittsburgh, PA).

### 2.2. Tissue Preparation

Before sectioning, all ITO slides were soaked in saturated sodium carbonate solution for at least 30 min. The slides were then washed with water and methanol and dried before mounting tissue sections. Frozen bovine lenses were sectioned to 10 μm thickness equatorially or axially using a CM 3050 Cryostat (LEICA CM 3050S, Leica Microsystems Inc., Bannockburn, IL). Sections were thaw-mounted on Indium-Tin-Oxide (ITO) Coated Slides (Delta Technologies, Loveland, CO). When sections on ITO slides were dry, the sections were washed sequentially as follows: 2% Triton X-100 in 50 mM ammonium formate for 5 min three times, 50 mM ammonium formate for 5 min, dried, water washed for 5 min twice, dried, PBS wash for 2 min three times, water wash for 2 min twice, dried, 0.002% formic acid wash for 2 min, water wash for 2 min, dried and 95% ethanol wash for 2 min twice, dried. Drying the sections between washes was done to maintain section morphology and all drying steps were done in a vacuum desiccator. To further remove crystallins from the sections for imaging the myristoylated filensin peptide, an additional wash with 2 M urea for 2 min was added after the PBS wash. Sections were desiccated before on-tissue digestion.

### 2.3. On-Tissue Digestion and MALDI Matrix Application

On-tissue digestion was performed as described previously (Judd et al., 2019). Briefly, the sections were sprayed with trypsin (15 ng/µL in 10% ACN in 100 mM ammonium bicarbonate, pH 8) using a TM Sprayer (HTX Technologies, Carrboro, NC, USA) modified with a syringe pump at 8 μL/min (Harvard Apparatus, Holliston, MA, USA). Trypsin was applied in eight passes with nozzle velocity of 750 mm/min at 30°C. The final trypsin concentration on tissue was 0.64 ng/mm^2^. Digestion was done in a sealed humidified petri dish at 37°C with 0.1 mL 100 mM ammonium bicarbonate overnight (16-18 hours). For combined LysC and trypsin digestion, the sections were sprayed with LysC (10 ng/µL in 10% ACN in 100 mM ammonium bicarbonate, pH 8, corresponding to final concentration of 0.43 ng/mm^2^) and incubated at 37°C for 5 hours as described for trypsin digestion. The sections were then sprayed with trypsin and digested by trypsin as described above. For GluC digestion, the section was sprayed with GluC (20 ng/uL 10% ACN in 100 mM ammonium bicarbonate, pH 8.0, corresponding to final 0.85 ng/mm^2^) and incubated as described for trypsin. After digestion, the sections were either sprayed with CHCA (5 mg/mL, 90% ACN, 0.1% TFA) or DHB (50 mg/mL, 90% ACN, 0.1% TFA) in 8 passes using the TM sprayer (85°C, 0.1 mL/min flow rate, 700 mm/min velocity, 2-mm track spacing, nitrogen carrier gas 9.5 psi). CHCA matrix was used for imaging tryptic peptides and LysC/trypsin or GluC digestion and DHB matrix was used for imaging the myristoylated filensin peptide.

### 2.4. MALDI Imaging Data Acquisition

Imaging experiments were performed using a Bruker SolariX 15T FTICR mass spectrometer (Bruker Daltonics, Billerica, MA, USA). Images were acquired using a Smartbeam II 2kHz Nd:YAG (355 nm) laser using the small laser setting (estimated spot size ∼50 μm) with 500 laser shots per pixel with a raster step size of 150 μm and a smart walk of 100 µm was enabled to increase sensitivity. The mass spectrometer was externally calibrated prior to analysis using a peptide mixture (Leu-enkephalin, Angiotensin II, Fibrinopeptide B, ACTH fragment (1-24) and insulin chain B). The mass spectrometer was operated in positive ion mode and data were collected from m/z 400 to 3500 at a mass resolution of ∼120,000 at m/z 1046.542. To achieve better sensitivity, continuous accumulation of selected ions (CASI) imaging (Fuchser et al., 2014) was used for imaging myristoylated filensin peptides with a Q1 mass of 471 (LysC + trypsin digestion) and 1146 (GluC digestion) and an isolation window of m/z 10. The raster step size was 200 μm for m/z 471 and 100 μm for m/z 1146. In CASI mode, 2000 laser shots were acquired per pixel.

### 2.5. Tissue extractions and LC-MS/MS analysis

To identify the peptide signals in MALDI imaging, peptides were generated from different regions of the lens using a liquid micro-enyzmatic digest and liquid extraction surface analysis (MicroLESA) from serial sections of lens tissue (Ryan et al., 2019b). Here, small volumes of trypsin (350pL, 0.075 μg/μL) were dispensed onto the lens at various positions (at normalized lens distance of 1, 0.89, 0.81, 0.77, 0.67 and 0). The trypsin spot size on tissue was estimated to be less than 200µm. In total, 15 individual drops were dispensed on tissue for each region of interest (ROI) (∼5.25 nL total). The tissue was incubated for 3 hours in a humidified petri dish at 37°C. Peptide extractions were completed using the TriVersa NanoMate (Advion, Inc., Ithaca, NY, USA) modified to include a glass capillary (LESAplus) for improved spatial resolution. LESA extraction used 2 μL, 20:80 ACN:H_2_O, 0.5% FA and two extracts per digest ROI were collected and combined prior to LC-MS/MS.

For LC-MS/MS analysis of the samples, tryptic peptides corresponding to half of the extract from each ROI were separated on a one-dimensional fused silica capillary column (200 mm x 100 µm) packed with Phenomenex Jupiter resin (3 µm mean particle size, 300 Å pore size) coupled with an UltiMate 3000 RSLCnano system (Thermo Scientific). An 82-minute gradient was performed, consisting of the following: 0-75 min, 2-45% ACN (0.1% formic acid); 75-79 min, 45-90% ACN (0.1% formic acid) balanced with 0.1% formic acid. The flow rate was 350 nL/min. The eluate was directly infused into a Q Exactive Plus instrument (Thermo Scientific, San Jose, CA) equipped with a nanoelectrospray ionization source. The data-dependent instrument method consisted of MS1 acquisition (R=70,000) from m/z 350-1600, using an MS AGC target value of 3e6, maximum ion time of 60 ms followed by up to 20 MS/MS scans (R=17,500) of the most abundant ions detected in the preceding MS scan. The MS2 AGC target value was set to 5e4, with a maximum ion time of 100 ms, HCD collision energy was set to 28, dynamic exclusion was set to 5 s.

### 2.6. Data Analysis

For MALDI imaging, ion images were generated using FlexImaging software (Version 4.1, Bruker Daltonics, Billerica, MA, USA). Each m/z signal was plotted ± 10 ppm and images were displayed using root-mean-square (RMS) normalization. Data were processed by FTMS processing 2.2.0 (Bruker Daltonics, Billerica, MA, USA) and recalibrated with four alpha crystallin peptides (αA 158-163, m/z 642.3933; αB 158-163, m/z 700.4352; αA 104-122, m/z 1090.4912; αA 146-157, m/z 1224.5967). The processed data were imported into SCiLS lab software (version 2017b, SCiLS, Bremen, Germany) and normalized by root-mean-square (RMS). Regional segmentation was performed using a bisecting k-means method and principal components were analyzed using pLSA with random initialization for 5 components. Peptide signals within each segmented region were identified based on LC-MS/MS data. A mass tolerance of 10 ppm was used when assigning imaged signals from LC-MS/MS data. FIJI software (https://fiji.sc) was used for measuring the distance and the normalized lens distance was defined as (r/a) where (a) represents the lens radius and (r) represents the distance from the lens core to the region of interest. To obtain lens radius (a), the diameter was measured using the average of measurements of ten different positions. To obtain (r), the distances from the capsule to the regions of interest were measured using the average of ten measurements at different positions and (r) was then calculated. Only regions where the capsule was clearly visible were used for measurements.

For analyzing LC-MS/MS data, the raw data were loaded in MaxQuant software (http://maxquant.org/, version 1.6.6.0) (Cox and Mann, 2008) and searched against a UniProt bovine database (downloaded 12/07/2016). The following search parameters were used: enzyme: trypsin, maximum miss cleavage: 2, precursor mass tolerance: 5 ppm, variable modifications: protein N-terminal acetylation and methionine oxidation. 1% PSM FDR and 1% protein FDR was used for protein identification.

## 3. Results

### 3.1. Preparation of lens sections

Lens tissue is filled with highly abundant crystallins. Due to ion suppression, low-abundance signals are challenging to detect using MALDI imaging without at least partial removal of lens crystallins. Multiple washing conditions were tested and the goal for optimization was to maintain the section morphology, retain lens intermediate filament proteins, and remove crystallins as much as possible. One step, found to be very helpful for retaining the section on slides during multiple steps of washing, was to soak ITO slides in saturated sodium carbonate solution for at least 30 minutes prior to tissue sectioning. Optimization of washing conditions resulted in a washing method that includes multiple steps of washing as described in the method section. To maintain section morphology and to avoid tissue detachment from the slide during washes, sections were dried between some washes; however, soluble proteins, once dried, may become insoluble and resistant to subsequent washes. Therefore, the first washes were done without a drying step and were found to be the most important for selectively removing crystallins. Triton X-100 in 50 mM ammonium formate was found to be a good choice for early washes to maintain tissue morphology even after multiple washes. Maintenance of section morphology is correlated with retention of lens intermediate filament proteins as shown in Figure 1. As shown in Figure 1, a brief water wash removed a significant amount of vimentin (Figure 1, lane B). Multiple water washes resulted in partial detachment of the section from the slide as indicated by weak protein bands on the gel (Figure 1, lane C). Inclusion of Triton X-100 in water helped retain intermediate filament proteins, especially vimentin, on the slide (Figure 1, lane D) and Triton X-100 in 50 mM ammonium formate further helped to retain lens intermediate filament proteins (Figure 1, lane E). After a section was dried multiple times, the section morphology can be easily maintained and the section was then washed with solutions in different pH to further remove unwanted proteins. After all aqueous washes, a 95% ethanol wash was used to remove lipids.

**Figure 1.**
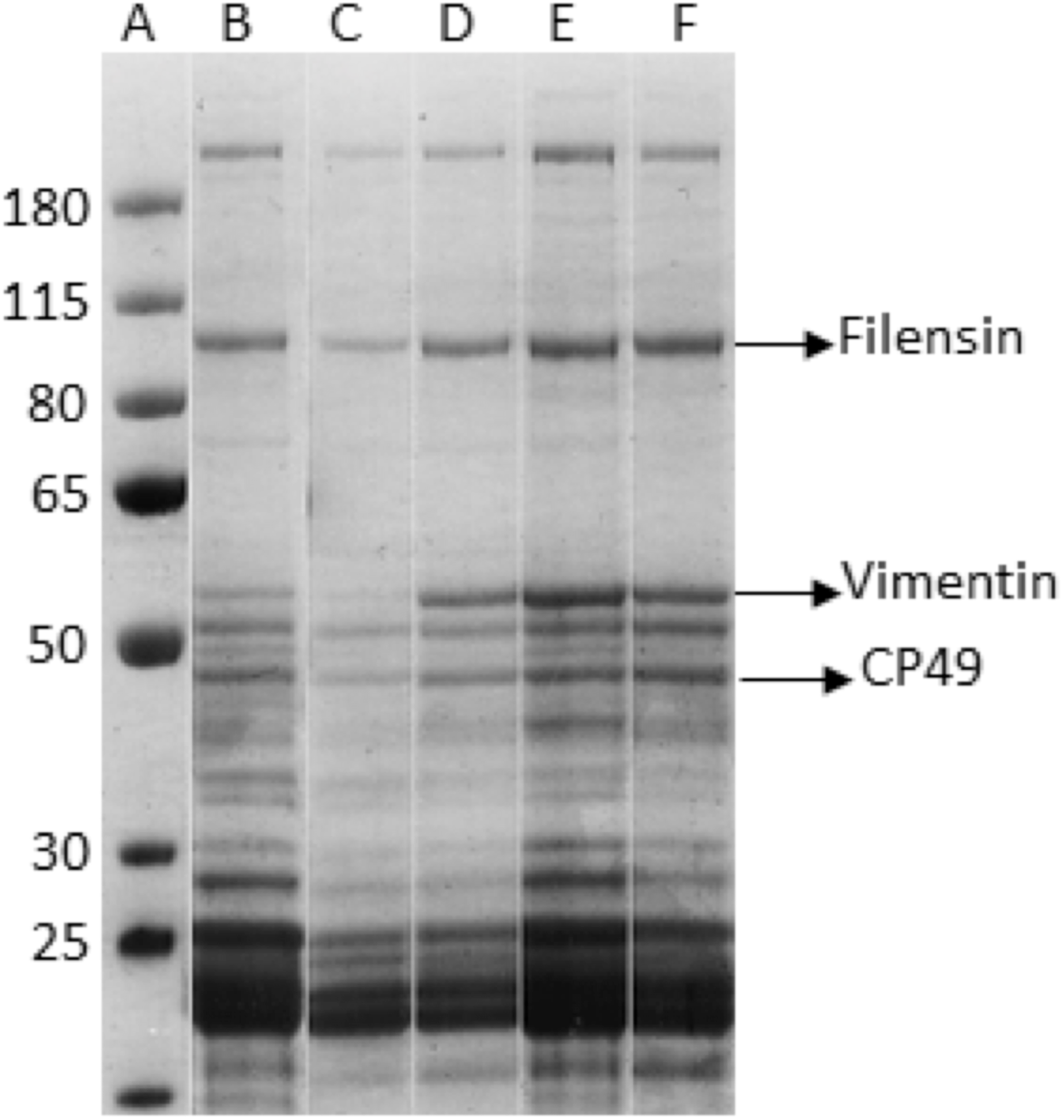
Optimization of initial washing steps to selectively remove crystallins. Lane A: molecular weight markers. The remaining lanes show the SDS-PAGE results (Coomassie Blue stain) of the proteins remaining on the slides in the cortex region after initial washing using different buffers. Lane B: the section was washed with water for 2 min.; Lane C: the section was washed with water for 2 min., twice; Lane D: the section was washed with 2% Triton X-100 in water for 5 min., twice, followed by water wash for 2 min.; Lane E: the section was washed with 2% Triton in 50 mM ammonium formate for 5 min., twice, followed by water wash for 2 min.; Lane F: the final section after all washing steps described in the method. The gel bands were assigned based on previously reported results (Carter, 1995, Wang, 2010).

### 3.2. Imaging of lens intermediate filament proteins

After optimization of washing conditions, enzyme digestion and data acquisition methods, rich peptide signals were observed from lens tissue. Overall, 1578 m/z features were detected at a S/N > 3 in the equatorial section and 1577 features were detected in the axial section. Identification of MALDI imaging signals was done by manual comparison between imaging data and LC-MS/MS data based on the accurate mass and specific regions where the signals were present. Therefore, identification was only attempted on the top 150 features that showed different distributions within the lens cortex as described in spatial segmentation section. Peptides from αA- and αB-crystallins still dominated the overall spectra, but multiple peptides from each of three lens intermediate filament proteins were detected. MALDI images of representative peptides are shown in Figure 2 (axial section) and Figure 3 (equatorial section). Peptide identification for signals shown in these figures were confirmed by their tandem mass spectra from LC-MS/MS analysis of regionally directed digestion and extraction. Strong vimentin signal was only detected in the superficial fiber cells and vimentin signal dramatically decreased at a normalized lens distance of approximately 0.9-0.91 from three different experiments (equatorial sections). Expression of lens beaded filament proteins started from the superficial fiber cells, but at the normalized lens distance of 0.9-0.91, both filensin and CP49 showed a dramatic signal increase. Although different peptides from the same protein showed different signal intensities, likely due to differences in ionization efficiencies, peptides from different protein regions (e.g. head, rod-like central portion, and tail) of each intermediate filament protein showed similar spatial distributions. Two peptides from each region were selected from vimentin and filensin and these signals are overlaid in Figure 2 (E, F) and 3 (E, F). Only one peptide from each region of CP49 was selected (Figure 2G and Figure 3G) because only one peptide from the tail region was detected due to the extremely short tail of CP49. For all three intermediate filament proteins, signals from different regions of the protein indicated by different peptides overlaid well. Therefore, no significant degradation of head or tail regions of these proteins was observed during fiber cell differentiation in bovine lens cortex. The sharply dropping signal of vimentin in the inner cortex seems to be due to the removal of all regions of vimentin since all peptide signals decreased in a similar pattern. Truncations in the head regions of filensin and CP49 reported previously (Wang et al., 2010) may only represent small fraction of these proteins. In addition, truncation in the head region of filensin results in increased water solubility (data not shown), therefore, this truncated form of filensin may have been removed during washes.

**Figure 2.**
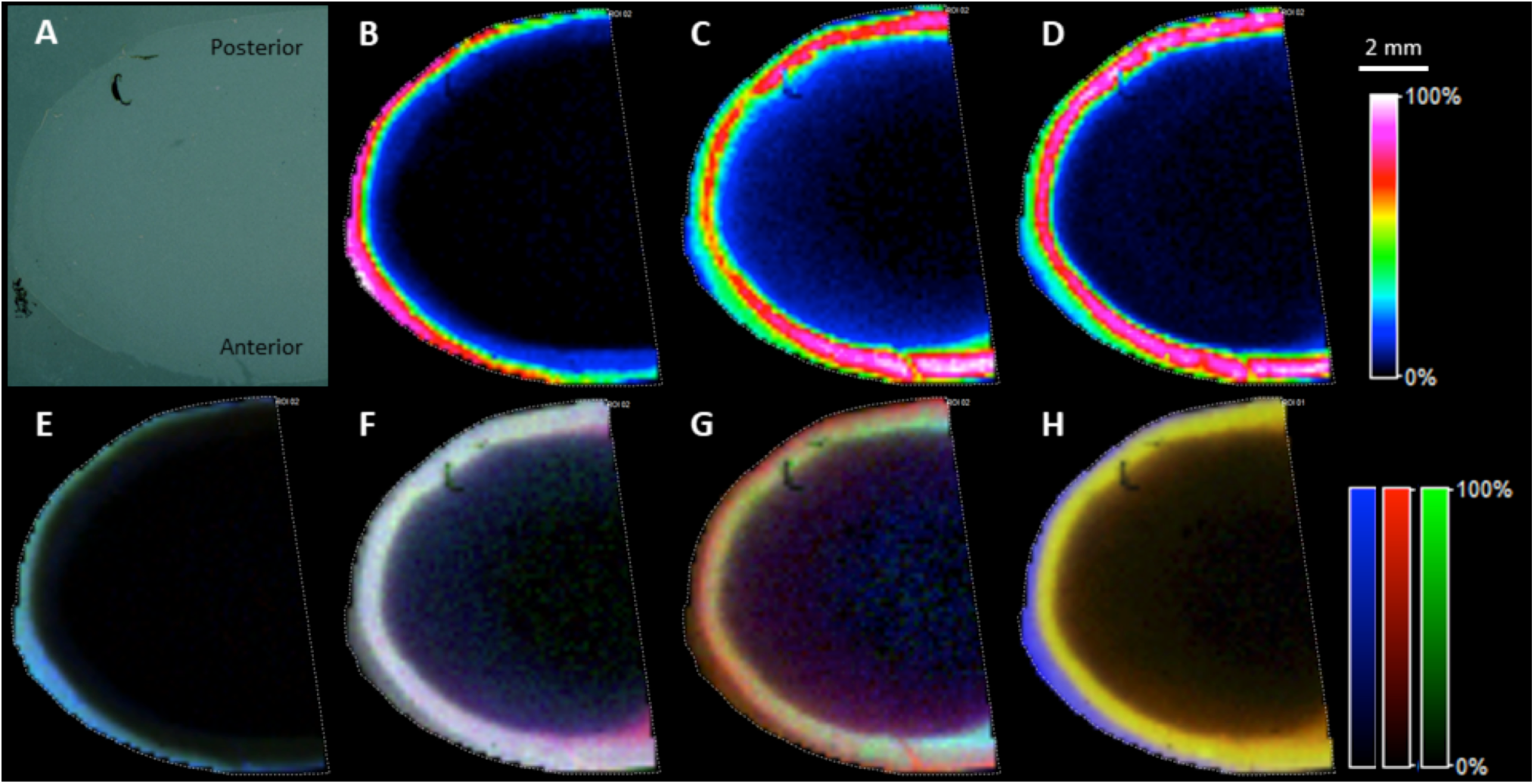
MALDI imaging of lens intermediate filament proteins in an axial lens section after on-tissue digestion. A: optical image; B: image for vimentin 37-50, m/z 1509.802; C: image for filensin 34-48, m/z 1451.7965; D: image for CP49 28-42, m/z 1456.753; E: overlay of signal from six vimentin peptides with peptides 29-36 and 37-50 from head region in green, peptides 105-113 and 295-304 from the central region in blue and peptides 411-424 and 446-450 from the tail region in red; F: overlay of signals from six filensin peptides with peptides 5-11 and 24-29 from the head region in green, peptides 59-63 and 77-89 from the central region in blue and peptides 351-361 and 677-693 from the tail region in red; G: overlay of signal from three CP49 peptides with peptide 6-21 from the head region in green, peptide 298-304 from the central region in blue and peptide 401-411 from the tail region in red. H: overlay six major peptide signals from vimentin in blue, filensin in red and CP49 in green. The six peptides from vimentin include peptides: 5-12, 29-36, 37-50, 51-64, 105-113 and 411-424. Six peptides from filensin include peptides: 5-11, 35-49, 59-63, 77-89, 351-361 and 677-693. Six peptides from CP49 include peptides: 29-43, 44-52, 77-89, 90-103, 128-137 and 401-411.

**Figure 3.**
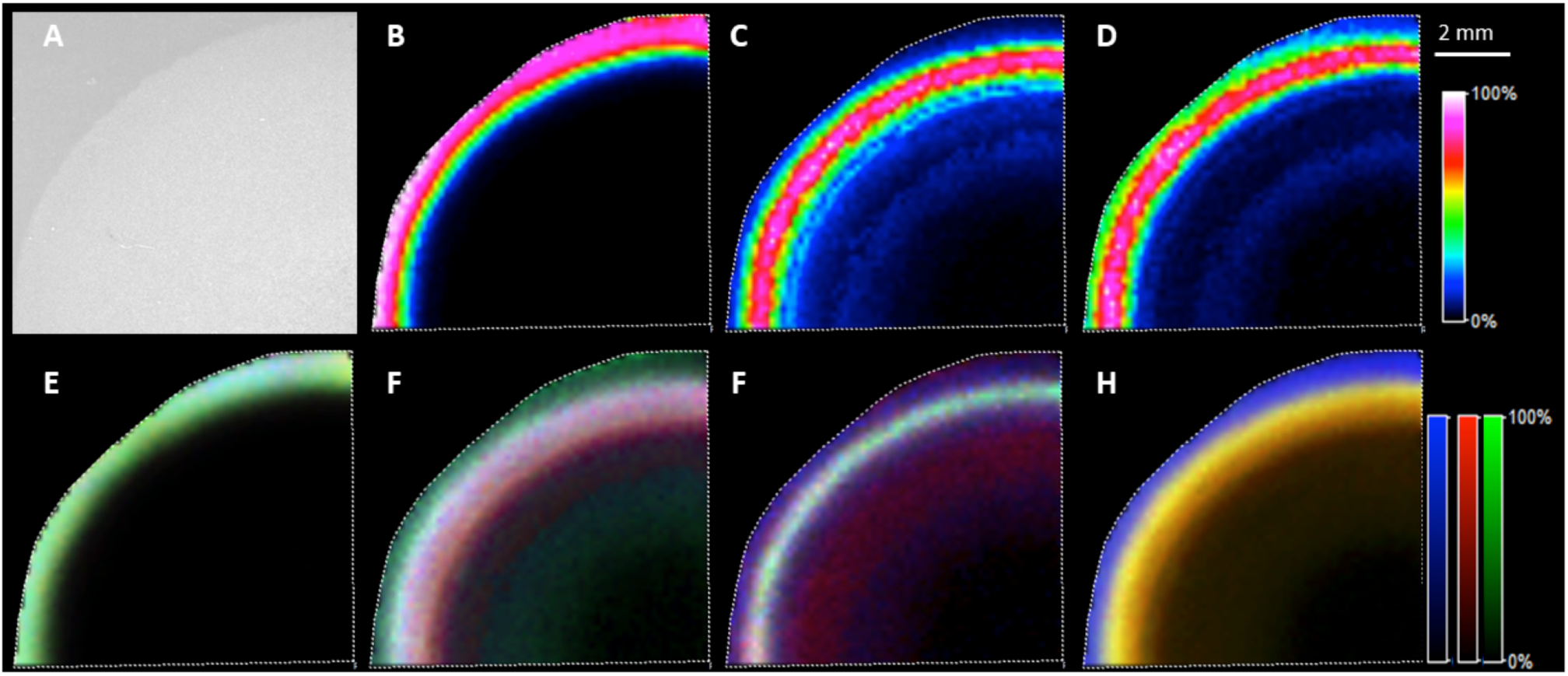
MALDI imaging of lens intermediate filament proteins in equatorial lens section after on-tissue digestion. A: optical image; B: image for vimentin 37-50, m/z 1509.802; C: image for filensin 34-48, m/z 1451.7965; D: image for CP49 28-42, m/z 1456.753; E: overlay of signal from six vimentin peptides with peptides 37-50 and 51-64 from head region in green, peptides 105-113 and 295-304 from the central region in blue and peptides 411-424 and 446-450 from the tail region in red; F: overlay of signals from six filensin peptides with peptides 24-29 and 30-34 from the head region in green, peptides 59-63 and 77-89 from the central region in blue and peptides 351-361 and 677-693 from the tail region in red; G: overlay of signal from three CP49 peptides with peptide 77-89 from the head region in green, peptide 192-199 from the central region in blue and peptide 401-411 from the tail region in red. H: overlay six major peptide signals from vimentin in blue, filensin in red and CP49 in green. The six peptides from vimentin include peptides: 14-28, 37-50, 51-64, 105-113, 158-163 and 411-424. Six peptides from filensin include peptides: 35-49, 50-56, 59-63, 77-89, 351-361 and 677-693. Six peptides from CP49 include peptides: 6-22, 29-43, 44-52, 77-89, 90-103 and 401-411.

To further confirm the sharp intermediate filament transition in an unbiased analysis, unsupervised spatial segmentation was done using SCiLS Lab software. The result is shown in Figure 4 (A, B). Datasets from both equatorial and axial sections, were segmented into two cortical regions similar to regions defined in Figure 2 and Figure 3. These two regions are named as outer cortex region and inner cortex region. The separation of these two regions occurred at a normalized lens distance of 0.9-0.91 measured using three IMS experiments from equatorial sections. This distance was the same as the normalized lens distance mentioned above that was calculated based on three peptide signals. The top 150 enriched features in each region were obtained from SCiLS and identification of the signals was done based on the LC-MS/MS data. The detailed data, including peptide identifications, for the enriched features in each region can be found in Supplemental Table 1 (equatorial section) and Supplemental Table 2 (axial section). The MALDI images of these enriched features can be found in Supplemental Figures 1-4. To identify these signals, the monoisotopic masses in imaging data were compared with precursor masses of the peptides identified in LC-MS/MS analysis. For MALDI data, sodium adducts of some peptides are common, therefore, sodium adducts were also considered. Manual *de novo* sequencing was also performed on some signals in LC-MS/MS that were not identified by automatic database searching. The manual *de novo* sequencing identified several nontryptic and posttranslationally modified peptides in vimentin. There are many features listed in the Supplemental Tables 1 and 2 that have not been identified. Manual inspection of LC-MS/MS data showed the presence of some of these signals based on accurate masses, but these signals have not been identified by a MaxQuant search. These signals may correspond to unanticipated posttranslationally modified peptides. Some signals in Supplemental Tables 1 and 2 were not present in the LC-MS/MS data suggesting they may represent non-peptide signals such as matrix adducts or unremoved lipids. It is also possible that peptides observed in one ionization mode, e.g. MALDI, do not appear in data using a different ionization mode, e.g. electrospray (Bodnar et al., 2003; Nayak et al., 2008). Summarized results for peptide identification can be found in Supplemental Table 3 and Figure 4C. 63-83% of the enriched signals in outer cortex and inner cortex regions were identified. Among the identified signals, 73% and 90% of these signals were from vimentin peptides in the outer cortex region of the axial and equatorial sections, respectively, whereas, in the inner cortex region, 87% and 91% of the identified signals were from beaded filament proteins in the axial and equatorial sections, respectively. Except for two unidentified signals in these two regions, there was no other signal found to be enriched in both regions as shown in Figure 4D. This result further supports the presence of a very sharp transition from vimentin to BFSPs at a normalized lens distance of 0.90-0.91 in the equatorial region. Results from principal component analysis shown in Figure 5 (A-B and F-G) also confirmed the well separated two regions in the lens cortex. Other than vimentin, filensin and CP49, multiple peptides from spectrins were also identified in both outer cortex and inner cortex regions, suggesting that this segmentation does not correlate well with the changing of spectrin expression; however, signals for spectrin peptides were weak, therefore, further study is needed to draw quantitative conclusions.

**Figure 4.**
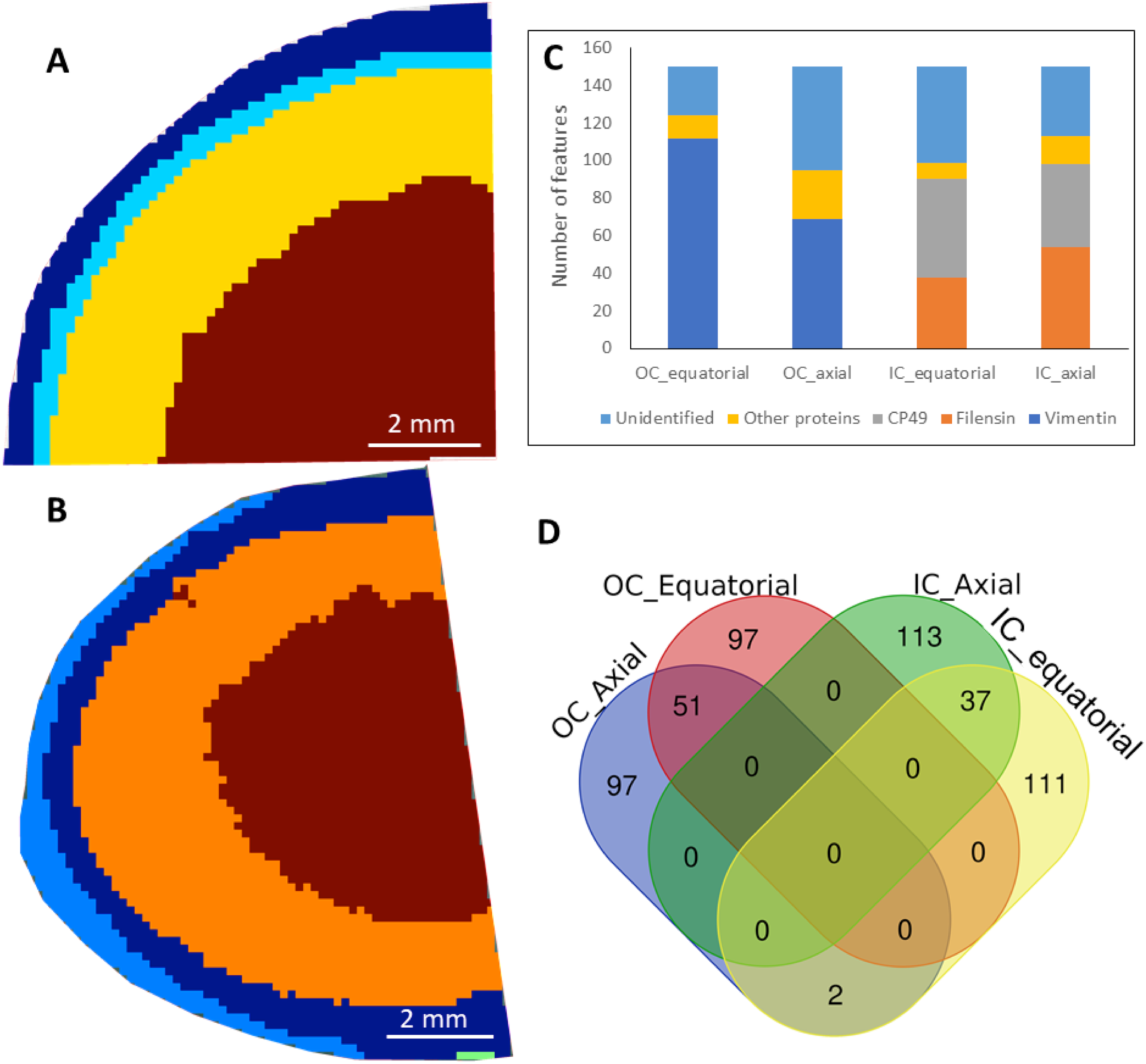
Spatial segmentation of lens IMS data determined using SCiLS software. Segmentation of the equatorial section image data is shown in A and segmentation for the axial section image data is shown in B. The two regions segmented in the cortex region were defined as outer cortex (OC) and inner cortex (IC) regions. Identification of the signals colocalized with each region is shown in C and vimentin was enriched in the outer cortex region and lens beaded filament proteins were enriched in the inner cortex region. D: Venn diagram shows signals enriched in OC and IC had virtually no overlap.

**Figure 5.**
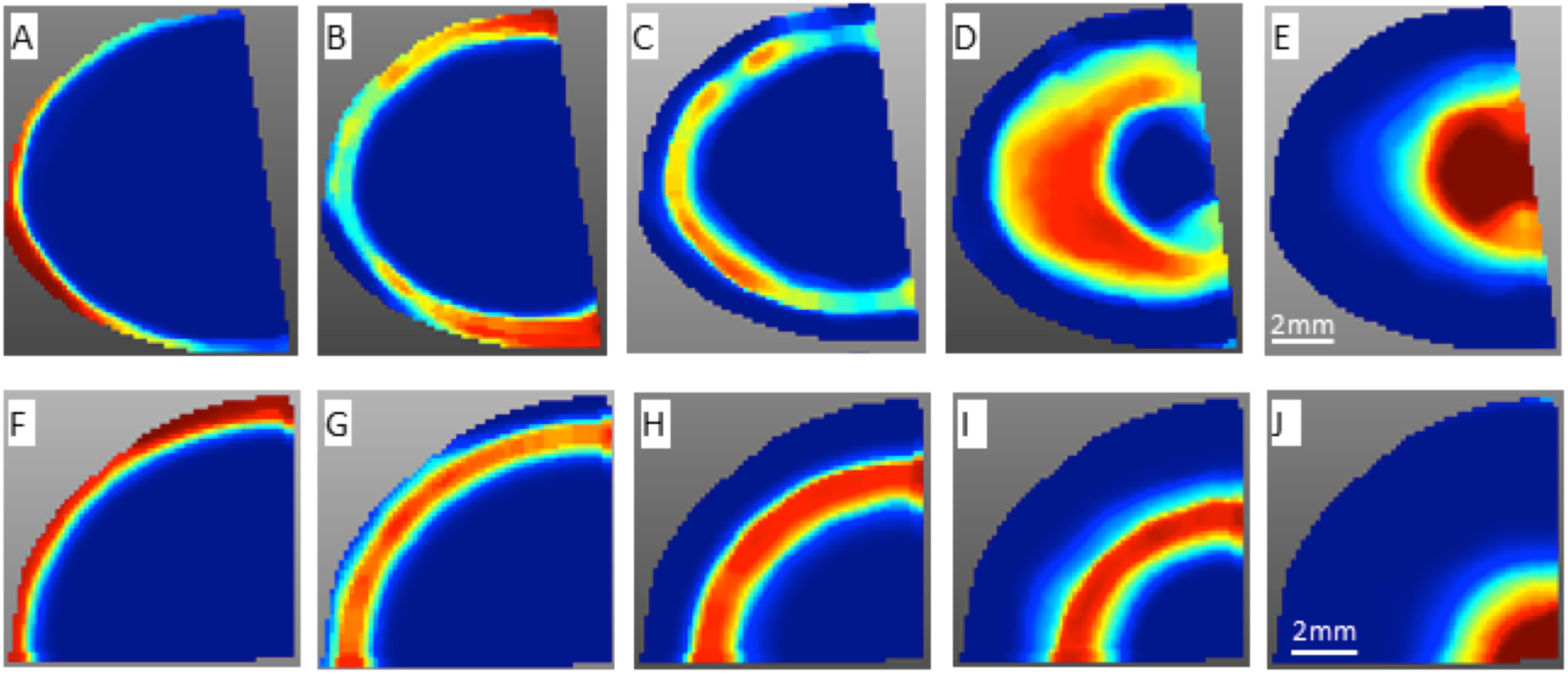
Principal component analysis (5 components) was performed using SCiLS. Components in the top row were from the dataset of the axial section and components in the bottom row were from dataset of the equatorial section. Components A-F and components B-G display similar outer cortex and inner cortex regions defined from spatial segmentation. Distinct regions in the nucleus were also identified.

Both spatial segmentation and principal component analysis of MALDI imaging data supported the presence of molecularly distinct regions in the lens nucleus (Figure 5). Enriched signals in the different nucleus regions have not been fully characterized, and peptides from αA- and αB-crystallins are among those enriched signals. Because the tissue section was extensively washed, the distribution of crystallins in our MALDI imaging data should only represent membrane associated crystallins. Based on the measured mass defect, many signals in the core of the lens are not derived from peptides.

The IMS results show that signals from all three intermediate filament proteins dropped dramatically in the lens nucleus and below the detection limit in the lens core by IMS. LC-MS/MS analysis of the extracts from different regions of the lens indicated that all three intermediate filament proteins were still detected in the lens core, but had weaker signal compared with the cortex region. Importantly, the MALDI imaging method used removes signals with low signal-to-noise ratio during data acquisition, therefore, low intensity signals do not appear in images of the lens core in IMS data indicating reduced signal of all three intermediate filament proteins. Signals for several nontryptic peptides were detected and all of these peptides are from the head region of vimentin (around residues 40-60) (Supplemental Table 1 and 2, Supplemental Figure 1 and 2). Images of these peptides showed they are present in the very young fiber cells; therefore, they are less likely due to age-related truncated vimentin. The generation of these peptides is likely to be due to either endogenous protease activity or to nonspecificity of trypsin. Another interesting finding is the detection of an N-terminal acetylated vimentin peptide corresponding to residues 51-64. The spatial localization of this acetylated peptide is similar to the other nonmodified vimentin peptides as shown in Supplemental Figure 1 (12F). The tandem mass spectrum for this peptide can be found in Supplemental Figure 5. The detection of N-acetylation of T51 suggests endogenous proteolysis may occur in the head region of vimentin around residue 51. Since N-acetylation protects against proteolysis (Jörnvall, 1975), this modification may indicate an attempt to prevent further protein degradation.

### 3.3. MALDI imaging of myristoylated filensin peptides

Previously, filensin was reported to undergo truncation and posttranslational myristoylation (Wang et al., 2010), but the region of the lens where the posttranslational myristoylation of filensin occurs has not been determined. Although it is challenging to obtain strong signal from fatty acylated peptides by MALDI, MALDI imaging has the advantage of providing spatial information. To improve sensitivity, continuous accumulation of selected ions (CASI) imaging on FT-ICR was performed. DHB was chosen as a matrix instead of more common matrix CHCA because DHB has been reported to improve signal intensity of fatty acylated peptides (Jagannadham and Nagaraj, 2005). In addition, trypsin alone was not efficient for digestion of the region of filensin that is myristoylated and a combination of LysC and trypsin digestion provided better efficiency of digestion. After optimization, MALDI imaging of the tryptic myristoylated filensin peptide 432-434 (GGK, m/z 471.3541) was obtained as shown in Figure 6A. The tryptic peptide 432-434 only has three residues and the same peptide can also be derived from brain acid soluble protein 1 (BASP1), that is also myristoylated at Gly 2 after Met 1 is removed. Furthermore, BASP1 is known to be present in the lens (Bassnett, 2009, Wang et al., 2013). Therefore, another proteolytic enzyme, GluC, was used to generate the filensin peptide 432-440 (GGKISKAFE, m/z 1146.7133) and this sequence is not conserved in other proteins based on a Protein Blast analysis of bovine proteins. The image of the myristoylated GluC peptide of filensin was obtained with better spatial resolution (100 µm) than the tryptic peptide 432-434 and the result is shown in Figure 6B. The identification of myristoylated 432-434 was confirmed by LC-MS/MS (data not shown). In addition, no BASP1 peptides were detected in microLESA experiment using the tissue section that had been washed using the same protocol as the imaging experiment. Therefore, we believe that BASP1 was removed during washing and the image of m/z 471.3541 is from myristoylated filensin 432-434. Overall the GluC peptide signal was weaker than tryptic myristoylated filensin peptide probably due to lower activity of GluC. Results from both images indicate that the posttranslational myristoylation of filensin starts at a normalized lens distance of 0.87. Similar to images for nonmodified filensin peptides, the signal for myristoylated peptides dramatically decreased in the lens core which is probably due to the reduced overall filensin abundance.

**Figure 6.**
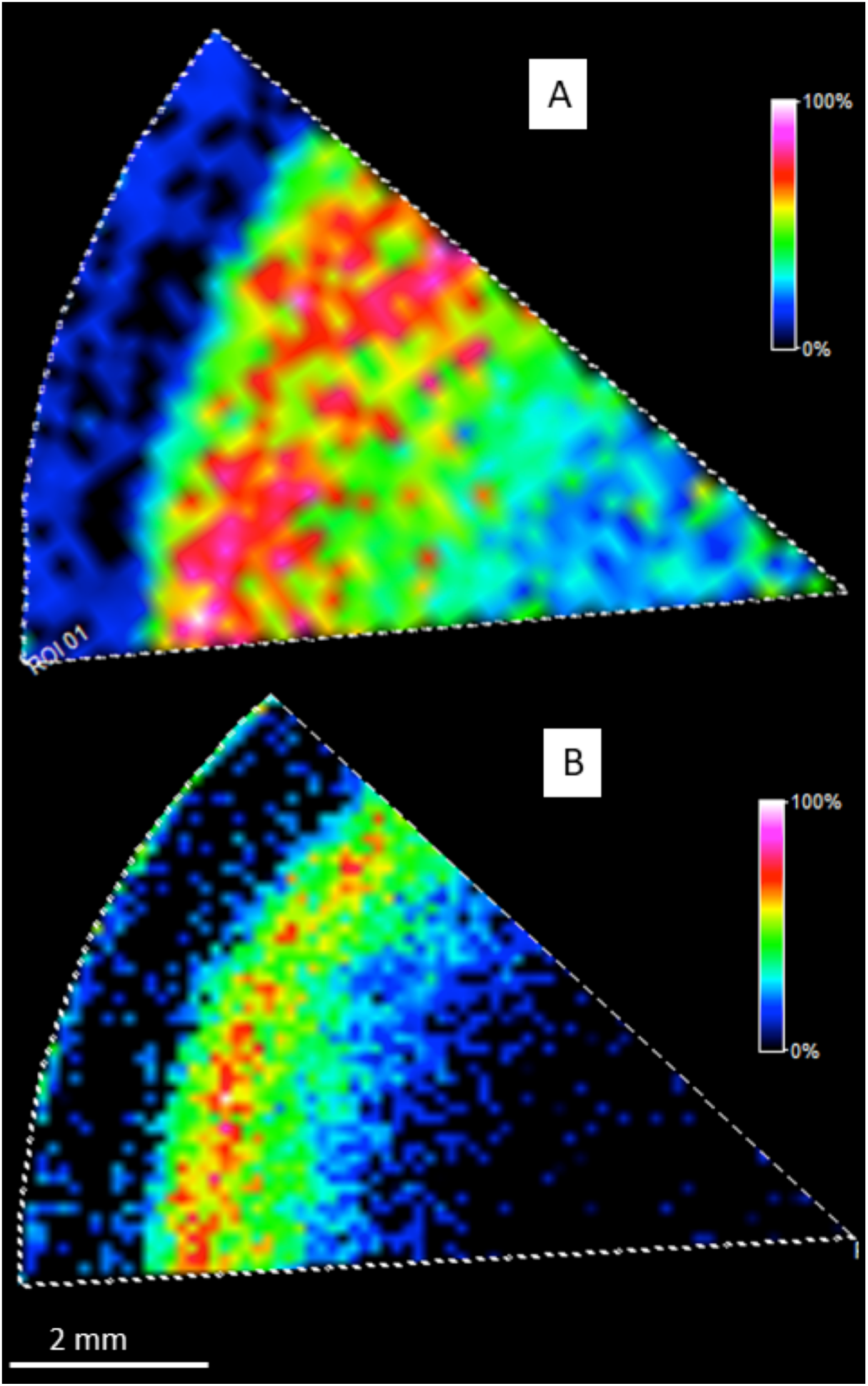
MALDI imaging of myristoylated filensin peptides. A: MALDI imaging of myristoylated 432-434 (m/z 471.3541±10 ppm) after a combination of LysC and Trypsin digestion. B: MALDI imaging of myristoylated 432-440 (m/z 1146.7133±10 ppm) after GluC digestion. Scale bar: 2 mm. Both results indicate that myristoylation starts at a normalized lens distance of 0.87.

## 4. Discussion

In this study, we developed a method for imaging cytoskeletal proteins that includes tissue washes, on-tissue digestion and data acquisition. This method should provide guidance for imaging cytoskeletal proteins in other tissues. Inclusion of Triton X-100 in the washing buffer is very important for retaining cytoskeletal proteins and for removing soluble proteins and some membrane proteins. With optimization of washing, this method is also expected to be useful for imaging other insoluble proteins in other tissues such as extracellular matrix proteins.

IMS data acquired using this novel sample preparation method show a switch from vimentin expression to beaded filament protein expression during fiber cell differentiation and maturation; a process that has been well documented in lens, albeit spatially ill-defined (Sandilands et al., 1995b; Blankenship et al., 2001). Our results show that before lens fiber cells completely lose vimentin signal, there is a region where vimentin signal significantly decreased, and this region spatially overlapped with the region where filensin/CP49 dramatically increased expression. Previously, an intermediate filament switch from vimentin to BFSPs was reported in the human lens outer cortex corresponding to the formation of the remodeling zone (Wenke et al., 2016). The results in this paper indicate that the intermediate filament switch is also present in bovine lens, even though the extreme morphologic changes present in human (Lim et al., 2009; Costello et al., 2013; Wenke et al., 2016) and other primate lenses (Costello et al., 2014) has not been reported in bovine lenses. The consistent detection of an intermediate filament switch in multiple species suggests that a conserved mechanism regulating intermediate filament networks exists between species. The switch from vimentin to BFSPs must be essential for maintenance of lens transparency since vimentin overexpression causes cataract (Capetanaki et al., 1989). Knockout studies indicate that beaded filaments are critical for maintaining interdigitations, differentiation state, and long-range stacking of lens fiber cells (Yoon et al., 2008). Therefore, increased expression of filensin and CP49 further into the lens cortex is critically important.

With fiber cell differentiation, vimentin was shown to disappear in the lens nucleus (Sandilands et al., 1995b; Blankenship et al., 2001) and filensin/CP49 signal also drastically decreased (Blankenship et al., 2001). This significant decrease of all three intermediate filament proteins in the lens center was also confirmed by our MALDI imaging data. Although absolute quantification by LC-MS/MS has not been done, signals for peptides from all three intermediate filament proteins can still be detected and are found to be low in lens core based on LC-MS/MS analysis of microLESA extracts. Therefore, all three intermediate filaments undergo a certain degree of degradation in the lens nucleus. Both filensin and CP49 undergo extensive truncation in the lens (Quinlan et al., 1999; Sandilands et al., 1995a). One of the major truncation sites in filensin is at residue D431 in bovine lens and D433 in human lens and truncation at this residue results in two fragments of filensin in the lens (Wang et al., 2010). We searched for signals corresponding to the C-terminus of the N-terminal fragment, unfortunately none were detected; however, the N-terminus of the C-terminal fragment was detected. MALDI images of the myristoylated N-terminal peptide indicate that proteolysis and posttranslational myristoylation occurred soon after the start of high expression of lens beaded filament proteins. Multiple roles of filensin in the lens have been predicted since it has an unusually long tail as an intermediate filament protein and this long tail is not required for its function as a co-assembling partner of CP49 (Masaki, 1997; Quinlan et al., 1999; Sandilands et al., 1995a). Interaction of filensin with the C-terminus of the lens major membrane protein, AQP0, has been reported (Nakazawa et al., 2011; Lindsey et al., 2006) and the direct interaction of filensin and AQP0 occurs in a similar lens region to where the formation of the filensin C-terminal fragment occurs. Truncation of filensin at D431 in bovine lens was previously found to start at a normalized distance of approximately 0.9 by analysis of dissected lens regions. Better spatial resolution provided by MALDI IMS shows that filensin myristoylation starts at a normalized distance of 0.87 and the results indicate that the formation of this C-terminal fragment starts early in the lens differentiation process. Therefore, lens cortical filensin undergoes multiple modifications and is expected to play an important role in fiber cell differentiation. Myristoylation of the C-terminal fragment is believed to be important to bring the C-terminal fragment to the membrane and interact with lens membrane proteins such as AQP0 (Sandilands et al., 1995a; Wang and Schey, 2017). This interaction has been reported to potentially drive AQP0 to redistribute in the membrane (Grey et al., 2009c) or to affect AQP0 permeability (Nakazawa et al., 2011).

The lens cortex region has been the focus of this study. Results from spatial segmentation and principal component analysis clearly show region-specific signals in the nucleus and these signals have not been thoroughly characterized. These signals are likely due to extensively modified peptides or, based on the observed mass defect, are non-peptidic in nature. Although IMS analysis of lens intermediate filament proteins was the focus of this study, the results provide information where distinct regions of protein expression and modification occur in the bovine lens. MALDI imaging-guided spatial proteomics and interactomics can then be performed to obtain more thorough understanding of protein-protein interaction and biological pathway changes associated with these distinct lens regions.

## Acknowledgements

This work was supported by NIH grants EY013462 and P30 EY008126.

## References

Anderson DM, Nye-Wood MG, Rose KL, Donaldson PJ, Grey AC, Schey KL. MALDI imaging mass spectrometry of β- and γ-crystallins in the ocular lens. J Mass Spectrom. 2020; 55:e4473;

Audette DS, Scheiblin DA, Duncan MK. The molecular mechanisms underlying lens fiber elongation. Exp Eye Res. 2017; 156:41–49.

Bassnett S, Wilmarth PA, David LL. The membrane proteome of the mouse lens fiber cell. Mol Vis. 2009; 15:2448–2463.

Blankenship TN, Hess JF, FitzGerald PG. Development- and differentiation-dependent reorganization of intermediate filaments in fiber cells. Invest Ophthalmol Vis Sci. 2001; 42: 735–742.

Bodnar WM, Blackburn RK, Krise JM, Moseley MA. Exploiting the complementary nature of LC/MALDI/MS/MS and LC/ESI/MS/MS for increased proteome coverage. J Am Soc Mass Spectrom. 2003; 14:971–979.

Buchberger AR, DeLaney K, Johnson J, Li L, Mass Spectrometry Imaging: A Review of Emerging Advancements and Future Insights. Anal Chem. 2018; 90:240–265.

Capetanaki Y, Smith S, Heath JP. Overexpression of the vimentin gene in transgenic mice inhibits normal lens cell differentiation. J Cell Biol. 1989; 109: 1653–1664.

Caprioli RM, Farmer TB, Gile J. Molecular imaging of biological samples: localization of peptides and proteins using MALDI-TOF MS. Anal Chem. 1997; 69:4751–4760.

Carter JM, Hutcheson AM, Quinlan RA, In vitro studies on the assembly properties of the lens beaded filament proteins: Co-assembly with α-crystallin but not with vimentin. Exp. Eye Res. 1995; 60: 181–192.

Costello MJ, Mohamed A, Gilliland KO, Fowler WC, Johnsen S. Ultrastructural analysis of the human lens fiber cell remodeling zone and the initiation of cellular compaction. Exp Eye Res. 2013; 116:411–418.

Costello MJ, Mohamed A, Gilliland KO, Wenke JL, Schey KL, Non-human primate lenses display the cortical remodeling zone, ARVO annual meeting Abstract, 2014.

Cox, J. and Mann, M. MaxQuant enables high peptide identification rates, individualized p.p.b.-range mass accuracies and proteome-wide protein quantification. Nat Biotechnol. 2008; 26, 1367–1372.

Deeley JM, Hankin JA, Friedrich MG, Murphy RC, Truscott RJ, Mitchell TW, Blanksby SJ. Sphingolipid distribution changes with age in the human lens. J Lipid Res. 2010; 51:2753–2760.

Ellis M, Alousi S, Lawniczak J, Maisel H, Welsh M, Studies on lens vimentin. Exp Eye Res. 1984; 38:195–202.

Farnsworth SC, Fu SCJ, Burke PA, Bahia I. Ultrastructure of rat eye lens fibers. Invest Ophthalmol. 1974; 13:274–279.

FitzGerald PG. Lens intermediate filaments. Exp Eye Res. 2009; 88:165–172.

Franck J, Longuespée R, Wisztorski M, Van Remoortere A, Van Zeijl R, Deelder A, Salzet M, McDonnell L, Fournier I. MALDI mass spectrometry imaging of proteins exceeding 30,000 daltons. Med Sci Monit. 2010; 16:BR293–299.

Fuchser J, Cornett S, Becker M. High resolution molecular imaging of pharmaceuticals at therapeutic levels. Application Note. 2014; # FTMS-37.

Grey AC and Schey KL. Distribution of Bovine and Rabbit Lens α-Crystallin Products by MALDI Imaging Mass Spectrometry, Mol. Vis. 2008; 14:171–179.

Grey AC and Schey KL. Age-related Changes in the Spatial Distribution of Human Lens α-Crystallin Products by MALDI Imaging Mass Spectrometry, Invest. Ophthalmol. Vis. Sci. 2009a; 50:4319–4329.

Grey AC, Chaurand P, Caprioli RM, Schey KL. MALDI imaging mass spectrometry of integral membrane proteins from ocular lens and retinal tissue. J. Proteom. Res., 2009b; 8:3278–3283.

Grey AC, Li L, Jacobs MD, Schey KL, Donaldson PJ. Differentiation-dependent modification and subcellular distribution of aquaporin-0 suggests multiple functional roles in the rat lens. Differentiation. 2009c; 77:70–83.

Groseclose, M R, Andersson, M, Hardesty, W M, Caprioli, R M, Identification of proteins directly from tissue: in situ tryptic digestions coupled with imaging mass spectrometry. J. Mass Spectrom. 2007; 42, 254–262.

Heeren RM, Smith DF, Stauber J, Kükrer-Kaletas B, MacAleese L. Imaging mass spectrometry: hype or hope? J Am Soc Mass Spectrom. 2009; 20:1006–1014.

Jagannadham MV, Nagaraj R. Detection of peptides covalently modified with multiple fatty acids by MALDI-TOF mass spectrometry. J Pept Res. 2005; 66:94–100.

Jörnvall H. Acetylation of Protein N-terminal amino groups structural observations on alpha-amino acetylated proteins. J Theor Biol. 1975; 55: 1–12.

Judd AM, Gutierrez DB, Moore JL, Patterson NH, Yang J, Romer CE, Norris JL, Caprioli RM. A recommended and verified procedure for in situ tryptic digestion of formalin-fixed paraffin-embedded tissues for analysis by matrix-assisted laser desorption/ionization imaging mass spectrometry. J Mass Spectrom. 2019; 54:716–727.

Kuwabara T. The maturation of the lens cell: a morphologic study. Exp Eye Res. 1975; 20:427–443.

Lemaire R, Desmons A, Tabet JC, Day R, Salzet M, Fournier I, Direct Analysis and MALDI Imaging of Formalin-Fixed, Paraffin-Embedded Tissue Sections, J. Proteome Res, 2007; 6. 1295–1305.

Lim JC, Walker KL, Sherwin T, Schey KL, Donaldson PJ. Confocal microscopy reveals zones of membrane remodeling in the outer cortex of the human lens. Invest Ophthalmol Vis Sci. 2009; 50:4304–4310.

Lindsey Rose KM, Gourdie RG, Prescott AR, Quinlan RA, Crouch RK, Schey KL. The C terminus of lens aquaporin 0 interacts with the cytoskeletal proteins filensin and CP49. Invest Ophthalmol Vis Sci. 2006; 47:1562–1570.

Lo WK, Harding CV. Square arrays and their role in ridge formation in human lens fibers. J Ultrastruct Res. 1984; 86: 228–245.

Lo WK, Biswas SK, Brako L, Shiels A, Gu S, Jiang JX. Aquaporin-0 targets interlocking domains to control the integrity and transparency of the eye lens. Invest Ophthalmol Vis Sci. 2014; 55:1202–1212.

Masaki S, Quinlan RA, Gene structure and sequence comparisons of the eye lens specific protein, filensin, from rat and mouse: implications for protein classification and assembly, Gene, 1997; 201: 11–20.

Nakazawa Y, Oka M, Furuki K, Mitsuishi A, Nakashima E, Takehana M. The effect of the interaction between aquaporin 0 (AQP0) and the filensin tail region on AQP0 water permeability. Mol Vis. 2011; 17:3191–3199.

Nayak R, Liu J, Sen AK, Knapp DR. Dual desorption electrospray ionization-laser desorption ionization mass spectrometry on a common nanoporous alumina platform for enhanced shotgun proteomic analysis. Anal Chem. 2008, 80:8840–8844.

Nicholl ID, Quinlan RA. Chaperone activity of alpha-crystallins modulates intermediate filament assembly. EMBO J. 1994; 13:945–953.

Nye-Wood MG, Spraggins JM, Caprioli RM, Schey KL, Donaldson PJ, Grey AC, Spatial distributions of glutathione and its endogenous conjugates in normal bovine lens and a model of lens aging. Exp Eye Res. 2017; 154:70–78.

Prentice BM, Ryan DJ, Van de Plas R, Caprioli RM, Spraggins JM. Enhanced ion transmission efficiency up to m/z 24000 for MALDI protein imaging mass spectrometry. Anal Chem. 2018;90:5090–5099.

Quinlan RA, Sandilands A, Procter JE, Prescott AR, Hutcheson AM, Dahm R, Gribbon C, Wallace P, Carter JM, The eye lens cytoskeleton. Eye. 1999; 13: 409–416.

Ryan DJ, Spraggins JM, Caprioli MR. Protein identification strategies in MALDI imaging mass spectrometry: a brief review. Curr Opin Chem Biol. 2019a;48:64–72.

Ryan DJ, Patterson NH, Putnam NE, Wilde AD, Weiss A, Perry WJ, Cassat JE, Skaar EP, Caprioli RM, Spraggins JM. MicroLESA: Integrating Autofluorescence Microscopy, In Situ Micro-Digestions, and Liquid Extraction Surface Analysis for High Spatial Resolution Targeted Proteomic Studies. Anal Chem. 2019b; 91:7578–7585.

Sandilands A, Prescott AR, Hutcheson AM, et al. Filensin is proteolytically processed during lens fiber cell differentiation by multiple independent pathways. Eur J Cell Biol. 1995a; 67:238–253.

Sandilands A, Prescott AR, Carter JM, Hutcheson AM, Quinlan RA, Richards J, FitzGerald PG. Vimentin and CP49/filensin form distinct networks in the lens which are independently modulated during lens fibre cell differentiation. J Cell Sci. 1995b; 108:1397–1406.

Schey KL, Gutierrez DB, Wang Z, Wei J, Grey AC. Novel fatty acid acylation of lens integral membrane protein aquaporin-0. Biochemistry. 2010; 49:9858–9865.

Seeley EH and Caprioli RM, Molecular imaging of proteins in tissues by mass spectrometry, Proc Natl Acad Sci U S A. 2008; 105: 18126–18131.

Seng JA, Nealon JR, Blanksby SJ, Mitchell TW. Distribution of Glycerophospholipids in the Adult Human Lens. Biomolecules. 2018; 8. pii: E156.

Song S, Landsbury A, Dahm R, Liu Y, Zhang Q, Quinlan RA. Functions of the intermediate filament cytoskeleton in the eye lens. J Clin Invest. 2009; 119:1837–1848.

Spivey EC, McMillen JC, Ryan DJ, Spraggins JM, Caprioli RM. Combining MALDI-2 and transmission geometry laser optics to achieve high sensitivity for ultra-high spatial resolution surface analysis. J Mass Spectrom. 2019; 54:366–370.

Spraggins JM, Rizzo DG, Moore JL, Noto MJ, Skaar EP, Caprioli RM. Next-generation technologies for spatial proteomics: integrating ultra-high speed MALDI-TOF and high mass resolution MALDI FTICR imaging mass spectrometry for protein analysis. Proteomics 2016;16:1678–1689.

Stella DR, Floyd KA, Grey AC, Renfrow MB, Schey KL, Barnes S. Tissue localization and solubilities of αA-crystallin and its numerous C-terminal truncation products in pre- and post-cataractous ICR/f rat lenses. Invest. Ophthalmol. Vis. Sci., 2010; 51:5153–5161.

Stoeckli M, Chaurand P, Hallahan DE, Caprioli RM. Imaging mass spectrometry: a new technology for the analysis of protein expression in mammalian tissues. Nat Med. 2001; 7:493–496.

Taylor VL, al-Ghoul KJ, Lane CW, Davis VA, Kuszak JR, Costello MJ. Morphology of the normal human lens. Invest Ophthalmol Vis Sci. 1996; 37:1396–1410.

Vérétout F, Delaye M, Tardieu A. Molecular basis of eye lens transparency. Osmotic pressure and X-ray analysis of alpha-crystallin solutions. J Mol Biol. 1989; 205:713–728.

Vidová V1, Pól J, Volny M, Novák P, Havlícek V, Wiedmer SK, Holopainen JM. Visualizing spatial lipid distribution in porcine lens by MALDI imaging high-resolution mass spectrometry. J Lipid Res. 2010; 51:2295–2302.

Vrensen G, Marie J, Veen H, Willekens B. Membrane architecture as a function of lens fiber maturation: A freeze fracture and scanning electron microscopic study in the human lens. Exp Eye Res. 1992; 54:433–446.

Wenke JL, Rose KL, Spraggins JM, Schey KL. MALDI Imaging Mass Specrometry Spatially Maps Age-Related Deamidation and Truncation of Human Lens Aquaporin-0. Invest. Ophthalmol. Vis. Sci., 2015; 56:7398–7405.

Wenke JL, McDonald WH, Schey KL. Spatially Directed Proteomics of the Human Lens Outer Cortex Reveals an Intermediate Filament Switch Associated With the Remodeling Zone. Invest Ophthalmol Vis Sci. 2016; 57:4108–4114.

Wang Z, Obidike JE, Schey KL, Posttranslational Modifications of the Bovine Lens Beaded Filament Proteins Filensin and CP49. Invest Ophthalmol Vis Sci. 2010; 51: 1565–1574.

Wang Z, Han J, David LL, Schey KL, Proteomics and phosphoproteomics analysis of human lens fiber cell membranes. Invest Ophthalmol Vis Sci. 2013; 54:1135–1143.

Wang Z, Schey KL. Identification of a direct Aquaporin-0 binding site in the lens-specific cytoskeletal protein filensin. Exp Eye Res. 2017; 159:23–29.

Yoon KH, Blankenship T, Shibata B, Fitzgerald PG. Resisting the effects of aging: a function for the fiber cell beaded filament. Invest Ophthalmol Vis Sci. 2008; 49: 1030–1036.

