## Supplementary Materials for "Localization of the Lens Intermediate Filament Switch by Imaging Mass Spectrometry"

Supplementary Table 1. Top 150 signal colocalized with outer cortex

| image | m/z | charge | Isotope number | Protein | peptide | theoretical MH+ | ppm |
|---|---|---|---|---|---|---|---|
| 1A | 510.11092 | 1 | 0 | | | | |
| 2A | 510.45197 | 1 | 0 | | | | |
| 3A | 519.28754 | 1 | 0 | Vimentin | SSAVR | 519.2885 | -1.848683343 |
| 4A | 547.28287 | 1 | 0 | | | | |
| 5A | 605.32458 | 1 | 0 | Vimentin | TVETR | 605.3253 | -1.189443098 |
| 6A | 606.32825 | 1 | 1 | Vimentin | TVETR | 606.32814 | 0.181419916 |
| 7A | 671.38278 | 1 | 0 | Vimentin | SGVPGVR | 671.3835 | -1.072412414 |
| 8A | 672.3862 | 1 | 1 | Vimentin | SGVPGVR | 672.38628 | -0.118979227 |
| 9A | 702.35168 | 1 | 0 | Vimentin | NNDALR | 702.352926 | -1.774036889 |
| 10A | 736.37527 | 2 | 0 | | | | |
| 11A | 736.87753 | 2 | 1 | | | | |
| 12A | 754.90814 | 2 | 0 | | | | |
| 13A | 755.40997 | 2 | 1 | | | | |
| 14A | 872.40936 | 1 | 0 | Vimentin | SVSSSSYR | 872.410836 | -1.691863442 |
| 15A | 873.41245 | 1 | 1 | Vimentin | SVSSSSYR | 873.41369 | -1.419716698 |
| 1B | 874.41574 | 1 | 2 | Vimentin | SVSSSSYR | 874.41618 | -0.503192885 |
| 2B | 905.42981 | 1 | 0 | Vimentin | GTNESLER | 905.432296 | -2.745649797 |
| 3B | 914.45587 | 1 | 0 | Vimentin | SYVTTSTR | 914.457786 | -2.09523067 |
| 4B | 915.45932 | 1 | 1 | Vimentin | SYVTTSTR | 915.46068 | -1.485590839 |
| 5B | 916.4628 | 1 | 2 | Vimentin | SYVTTSTR | 916.46323 | -0.469195038 |
| 6B | 925.39865 | 1 | 0 | Vimentin | QESNEYR | 925.400996 | -2.53511722 |
| 7B | 976.44568 | 1 | 0 | Actin, cytoplasmic 1/2 | AGFAGDDAPR | 976.448286 | -2.668856136 |
| 8B | 1028.51 | 1 | 0 | Vimentin | SVSSSSYRR | 1028.511976 | -1.911499376 |
| 9B | 1093.5259 | 1 | 0 | Vimentin | FADLSEAANR | 1093.527276 | -1.2217345 |
| 10B | 1094.5294 | 1 | 1 | Vimentin | FADLSEAANR | 1094.53013 | -0.648680178 |
| 11B | 1115.5067 | 1 | 0 | Vimentin | FADLSEAANR+Na | 1115.5091 | -2.142519501 |
| 12B | 1115.5676 | 1 | 0 | Vimentin | VELQELNDR | 1115.569176 | -1.385839653 |
| 13B | 1116.5724 | 1 | 1 | Vimentin | VELQELNDR | 1116.572 | 0.349283342 |
| 14B | 1189.5806 | 1 | 0 | Spectrin beta, non-erythrocytic 1 | LVSDGNINSDR | 1189.580776 | -0.17317025 |
| 15B | 1198.7031 | 1 | 0 | Actin, cytoplasmic 1/2 | AVFPSIVGRPR | 1198.705476 | -2.015507603 |
| 1C | 1199.7091 | 1 | 1 | Actin, cytoplasmic 1/2 | AVFPSIVGRPR | 1199.70835 | 0.583475142 |
| 2C | 1200.7119 | 1 | 2 | Actin, cytoplasmic 1/2 | AVFPSIVGRPR | 1200.71105 | 0.716242263 |
| 3C | 1203.591 | 1 | 0 | Spectrin beta, non-erythrocytic 1 | DQNTVETLQR | 1203.5964 | -4.528095963 |
| 4C | 1245.6889 | 1 | 0 | Vimentin | SLGSALRPTTSR (semitryptic) | 1245.690976 | -1.666544946 |
| 5C | 1246.6931 | 1 | 1 | Vimentin | SLGSALRPTTSR (semitryptic) | 1246.69376 | -0.513357828 |
| 6C | 1270.5644 | 1 | 0 | Vimentin | LGDLYEEEMR+Oxidation | 1270.562 | 1.888927892 |
| 7C | 1271.5649 | 1 | 1 | Vimentin | LGDLYEEEMR+Oxidation | 1271.56492 | -0.031457301 |
| 8C | 1278.5961 | 1 | 0 | Vimentin | EEAESTLQSFR(dehydration on N-te | 1278.5961 | 0.023463234 |
| 9C | 1296.6063 | 1 | 0 | Vimentin | EEAESTLQSFR | 1296.606676 | -0.320837466 |
| 10C | 1297.6096 | 1 | 1 | Vimentin | EEAESTLQSFR | 1297.60954 | 0.061651828 |
| 11C | 1302.6475 | 1 | 0 | Spectrin alpha, non-erythrocytic 1 | VLETAEDIQER | 1302.653576 | -4.656648638 |
| 12C | 1310.6329 | 1 | 0 | Spectrin beta, non-erythrocytic 1 | ALVADSHPESER | 1310.633476 | -0.416592442 |
| 13C | 1311.6377 | 1 | 1 | Spectrin beta, non-erythrocytic 1 | ALVADSHPESER | 1311.63637 | 1.01400055 |
| 14C | 1324.6948 | 1 | 0 | Spectrin alpha, non-erythrocytic 1 | SQLLGSAHEVQR | 1324.6968 | -1.494681651 |
| 15C | 1352.7147 | 1 | 0 | | | | |
| 1D | 1353.7002 | 1 | 0 | Vimentin | TYSLGSALRPTTS(semitryptic) | 1353.7009 | -0.517100934 |
| 2D | 1354.7042 | 1 | 1 | Vimentin | TYSLGSALRPTTS(semitryptic) | 1354.70378 | 0.280504126 |
| 3D | 1355.7095 | 1 | 2 | Vimentin | TYSLGSALRPTTS(semitryptic) | 1355.70647 | 2.257125763 |
| 4D | 1377.6857 | 1 | 0 | | | | |
| 5D | 1408.7548 | 1 | 0 | | | | |
| 6D | 1415.6668 | 1 | 0 | | | | |
| 7D | 1416.6653 | 1 | 1 | | | | |
| 8D | 1454.7261 | 1 | 0 | | | | |
| 9D | 1455.737 | 1 | 1 | | | | |
| 10D | 1460.7323 | 1 | 0 | | | | |
| 11D | 1461.736 | 1 | 1 | | | | |
| 12D | 1465.7767 | 1 | 0 | Spondin-1 | AQWPAWQPLNVR | 1465.769876 | 4.621462148 |
| 13D | 1466.7883 | 1 | 0 | | | | |
| 14D | 1467.7819 | 1 | 0 | Tubulin beta | FIGNSTAIQELFK | 1467.784276 | -1.646018451 |
| 15D | 1468.7853 | 1 | 1 | Tubulin beta | FIGNSTAIQELFK | 1468.78718 | -1.252734246 |
| 1E | 1472.7376 | 1 | 0 | Vimentin | TLYTSSPGGVYATR | 1472.737976 | -0.255306787 |
| 2E | 1473.7099 | 1 | 0 | Monocarboxylate transporter 1 | DEETNVDVAEKPK | 1473.706776 | 2.119824684 |
| 3E | 1473.7429 | 1 | 1 | Vimentin | TLYTSSPGGVYATR | 1473.74092 | 1.357090634 |
| 4E | 1474.746 | 1 | 2 | Vimentin | TLYTSSPGGVYATR | 1474.74365 | 1.573154765 |
| 5E | 1475.7504 | 1 | 3 | Vimentin | TLYTSSPGGVYATR | 1475.74629 | 2.764702868 |
| 6E | 1487.7612 | 1 | 0 | Reticulon | DTLAPDEVSALTQK | 1487.758776 | 1.609131829 |
| 7E | 1490.7144 | 1 | 0 | Fructose-bisphosphate aldolase | LQSIGTENTEENR | 1490.708176 | 4.188613238 |
| 8E | 1491.7009 | 1 | 0 | | | | |
| 9E | 1491.7907 | 1 | 0 | | | | |
| 10E | 1492.7065 | 1 | 1 | | | | |
| 11E | 1494.7245 | 1 | 0 | Vimentin | TLYTSSPGGVYATR+Na | 1494.7198 | 3.137711831 |
| 12E | 1494.7836 | 1 | 0 | Ankyrin-2 | SGHDQVVELLLER | 1494.7911 | -5.037493199 |
| 13E | 1495.7267 | 1 | 1 | Vimentin | TLYTSSPGGVYATR+Na | 1495.722863 | 2.551943341 |
| 14E | 1496.7303 | 1 | 2 | Vimentin | TLYTSSPGGVYATR+Na | 1496.725593 | 3.138183794 |

| | | | | | | | |
|---|---|---|---|---|---|---|---|
| 15E | 1507.7897 | 1 | 0 | | | | |
| 1F | 1508.7312 | 1 | 0 | Vimentin | MFGGPGTASRPSSTR | 1508.727476 | 2.468305283 |
| 2F | 1509.7357 | 1 | 1 | Vimentin | MFGGPGTASRPSSTR | 1509.73021 | 3.649658703 |
| 3F | 1509.7975 | 1 | 0 | Vimentin | TYSLGSALRPTTSR | 1509.801976 | -2.971250582 |
| 4F | 1510.734 | 1 | 0 | | | | |
| 5F | 1510.7717 | 1 | 0 | | | | |
| 6F | 1510.8049 | 1 | 1 | Vimentin | TYSLGSALRPTTSR | 1510.80483 | 0.066189886 |
| 7F | 1510.8306 | 1 | 0 | | | | |
| 8F | 1511.7366 | 1 | 0 | Spectrin alpha, non-erythrocytic 1 | MQHNLEQQIQAR+oxidation | 1511.7383 | -1.104688556 |
| 9F | 1511.8062 | 1 | 1 | | | | |
| 10F | 1512.8117 | 1 | 0 | Guanine nucleotide-binding protein s | DALLVGVPAGSNPFR | 1512.816876 | -3.414821769 |
| 11F | 1513.8156 | 1 | 1 | Guanine nucleotide-binding protein s | DALLVGVPAGSNPFR | 1513.81981 | -2.774438524 |
| 12F | 1514.7495 | 1 | 0 | Vimentin | acetyl-TLYTSSPGGVYATR | 1514.7485 | 0.666777356 |
| 13F | 1515.7195 | 1 | 0 | | | | |
| 14F | 1515.7545 | 1 | 1 | Vimentin | acetyl-TLYTSSPGGVYATR | 1515.7515 | 1.9528267 |
| 15F | 1521.803 | 1 | 0 | Spectrin beta, non-erythrocytic 1 | HALVEADIGIQAER | 1521.802 | 0.643973395 |
| 1G | 1522.8066 | 1 | 1 | Spectrin beta, non-erythrocytic 1 | HALVEADIGIQAER | 1522.80484 | 1.142628362 |
| 2G | 1524.7225 | 1 | 0 | Vimentin | MFGGPGTASRPSSTR+Oxidation | 1524.7224 | 0.045909996 |
| 3G | 1525.7284 | 1 | 1 | Vimentin | MFGGPGTASRPSSTR+Oxidation | 1525.72512 | 2.143243207 |
| 4G | 1525.8007 | 1 | 0 | | | | |
| 5G | 1526.7325 | 1 | 2 | Vimentin | MFGGPGTASRPSSTR+Oxidation | 1526.72669 | 3.831727079 |
| 6G | 1527.7365 | 1 | 3 | Vimentin | MFGGPGTASRPSSTR+Oxidation | 1527.72813 | 5.485269162 |
| 7G | 1530.7128 | 1 | 0 | Vimentin | MFGGPGTASRPSSTR+Na | 1530.7092 | 2.371449783 |
| 8G | 1531.7158 | 1 | 1 | Vimentin | MFGGPGTASRPSSTR+Na | 1531.712153 | 2.394052951 |
| 9G | 1531.7887 | 1 | 0 | Vimentin | TYSLGSALRPTTSR+Na | 1531.7838 | 3.198884856 |
| 10G | 1532.7884 | 1 | 1 | Vimentin | TYSLGSALRPTTSR+Na | 1532.786773 | 1.054941254 |
| 11G | 1546.7045 | 1 | 0 | Vimentin | MFGGPGTASRPSSTR+Oxidation+N | 1546.7042 | 0.174564729 |
| 12G | 1547.708 | 1 | 1 | Vimentin | MFGGPGTASRPSSTR+Oxidation+N | 1547.707063 | 0.573105868 |
| 13G | 1551.8176 | 1 | 0 | | | | |
| 14G | 1552.8214 | 1 | 1 | | | | |
| 15G | 1553.8216 | 1 | 1 | | | | |
| 1H | 1570.8976 | 1 | 0 | Vimentin | ISLPLPNFSSLNLR | 1570.895176 | 1.53033763 |
| 2H | 1571.7269 | 1 | 0 | | | | |
| 3H | 1571.9 | 1 | 1 | Vimentin | ISLPLPNFSSLNLR | 1571.89808 | 1.23417671 |
| 4H | 1572.7271 | 1 | 1 | | | | |
| 5H | 1572.9005 | 1 | 2 | Vimentin | ISLPLPNFSSLNLR | 1572.90086 | -0.222518792 |
| 6H | 1573.9022 | 1 | 3 | Vimentin | ISLPLPNFSSLNLR | 1573.90355 | -0.883154498 |
| 7H | 1592.8815 | 1 | 0 | Vimentin | ISLPLPNFSSLNLR+Na | 1592.877 | 2.806243043 |
| 8H | 1593.8853 | 1 | 1 | Vimentin | ISLPLPNFSSLNLR+Na | 1593.880023 | 3.27941873 |
| 9H | 1628.8439 | 1 | 0 | Vimentin | RTLYTSSPGGVYATR(semitryptic) | 1628.839076 | 2.94320051 |
| 10H | 1629.8479 | 1 | 1 | Vimentin | RTLYTSSPGGVYATR(semitryptic) | 1629.84197 | 3.638389555 |
| 11H | 1637.8998 | 1 | 0 | | | | |
| 12H | 1652.8312 | 1 | 0 | | | | |
| 13H | 1665.9078 | 1 | 0 | | | | |
| 14H | 1666.9121 | 1 | 1 | | | | |
| 15H | 1668.8507 | 1 | 0 | Vimentin | ETNLDSLPLVDTHSK | 1668.8439 | 4.080669258 |
| 1I | 1669.853 | 1 | 1 | Vimentin | ETNLDSLPLVDTHSK | 1669.84683 | 3.712915394 |
| 2I | 1672.8209 | 1 | 0 | | | | |
| 3I | 1673.8262 | 1 | 0 | Peptidyl-prolyl cis-trans isomerase Fk | DHLVTAYNHLFESK | 1673.828176 | -1.162604399 |
| 4I | 1674.8271 | 1 | 1 | Peptidyl-prolyl cis-trans isomerase Fk | DHLVTAYNHLFESK | 1674.83112 | -2.406212753 |
| 5I | 1690.8337 | 1 | 0 | Vimentin | ETNLDSLPLVDTHSK+Na | 1690.8257 | 4.719587595 |
| 6I | 1691.8342 | 1 | 1 | Vimentin | ETNLDSLPLVDTHSK+Na | 1691.828773 | 3.225503719 |
| 7I | 1701.9069 | 1 | 0 | Tubulin alpha-1D chain | AVFVDLEPTVIDEVR | 1701.905776 | 0.636933028 |
| 8I | 1706.7812 | 1 | 0 | | | | |
| 9I | 1754.8346 | 1 | 0 | | | | |
| 10I | 1755.8409 | 1 | 0 | | | | |
| 11I | 1776.8668 | 1 | 0 | Vimentin | FADLSEAANRNNDALR | 1776.862376 | 2.501037818 |
| 12I | 1777.8715 | 1 | 1 | Vimentin | FADLSEAANRNNDALR | 1777.86515 | 3.549200568 |
| 13I | 1806.9299 | 1 | 0 | Ankynin-2 | IENPNSLQDQSHALLK | 1806.934276 | -2.438384206 |
| 14I | 1807.9342 | 1 | 0 | | | | |
| 15I | 1818.7938 | 1 | 0 | | | | |
| 1J | 1819.7786 | 1 | 0 | | | | |
| 2J | 1820.7819 | 1 | 0 | | | | |
| 3J | 1830.7238 | 1 | 1 | | | | |
| 4J | 1836.8044 | 1 | 0 | Vimentin | DGQVINETSQHHDDLE | 1836.799476 | 2.702526903 |
| 5J | 1837.7954 | 1 | 0 | | | | |
| 6J | 1838.8008 | 1 | 1 | | | | |
| 7J | 1839.8029 | 1 | 0 | | | | |
| 8J | 1858.7777 | 1 | 0 | Vimentin | DGQVINETSQHHDDLE+Na | 1858.7813 | -1.931372992 |
| 9J | 1860.7718 | 1 | 0 | | | | |
| 10J | 1861.7727 | 1 | 1 | | | | |
| 11J | 1874.7421 | 1 | 0 | | | | |
| 12J | 1875.7349 | 1 | 0 | | | | |
| 13J | 1876.7404 | 1 | 1 | | | | |
| 14J | 1896.7313 | 1 | 0 | | | | |
| 15J | 1898.7172 | 1 | 0 | | | | |

Supplementary Table 2. Top 150 signal in outer cortex

| image | m/z | charge | Isotope number | Protein | peptide | theoretical MH+ | ppm |
|---|---|---|---|---|---|---|---|
| 1A | 519.28796 | 1 | 0 | Vimentin | SSAVR | 519.2885 | -1.039884 |
| 2A | 520.29126 | 1 | 1 | Vimentin | SSAVR | 520.29126 | 0 |
| 3A | 547.28308 | 1 | 0 | | | | |
| 4A | 605.32513 | 1 | 0 | Vimentin | TVETR | 605.3253 | -0.280841 |
| 5A | 606.328 | 1 | 1 | Vimentin | TVETR | 606.32814 | -0.230898 |
| 6A | 631.34033 | 1 | 0 | Vimentin | VEVER | 631.341 | -1.061233 |
| 7A | 632.34375 | 1 | 1 | Vimentin | VEVER | 632.34383 | -0.126513 |
| 8A | 645.35596 | 1 | 0 | Vimentin | LDLER | 645.3566 | -0.9917 |
| 9A | 646.35919 | 1 | 1 | Vimentin | LDLER | 646.3595 | -0.479609 |
| 10A | 671.38293 | 1 | 0 | Vimentin | SGVPGVR | 671.3835 | -0.848993 |
| 11A | 672.38599 | 1 | 1 | Vimentin | SGVPGVR | 672.38628 | -0.4313 |
| 12A | 673.38934 | 1 | 2 | Vimentin | SGVPGVR | 673.38875 | 0.8761655 |
| 13A | 693.36438 | 1 | 0 | Vimentin | SGVPGVR+Na | 693.3653 | -1.326862 |
| 14A | 694.36823 | 1 | 1 | Vimentin | SGVPGVR+Na | 694.368223 | 0.0100811 |
| 15A | 702.35248 | 1 | 0 | Vimentin | NNDALR | 702.352926 | -0.635008 |
| 1B | 703.33624 | 1 | 0 | Vimentin | NNDALR+deamidation | 703.3369 | -0.938384 |
| 2B | 703.35553 | 1 | 1 | Vimentin | NNDALR | 703.35563 | -0.142176 |
| 3B | 714.35229 | 1 | 0 | | | | |
| 4B | 724.33423 | 1 | 0 | Vimentin | NNDALR+Na | 724.3347 | -0.648871 |
| 5B | 754.91711 | 2 | 0 | | | | |
| 6B | 872.41125 | 1 | 0 | Vimentin | SVSSSSYR | 872.410836 | 0.4745471 |
| 7B | 873.41394 | 1 | 1 | Vimentin | SVSSSSYR | 873.41369 | 0.2862332 |
| 8B | 874.41675 | 1 | 2 | Vimentin | SVSSSSYR | 874.41618 | 0.6518635 |
| 9B | 894.39404 | 1 | 0 | Vimentin | SVSSSSYR+Na | 894.3926 | 1.6100312 |
| 10B | 905.43213 | 1 | 0 | Vimentin | GTNESLER | 905.432296 | -0.183338 |
| 11B | 906.43549 | 1 | 1 | Vimentin | GTNESLER | 906.43511 | 0.4192247 |
| 12B | 908.37451 | 1 | 0 | | | | |
| 13B | 914.45795 | 1 | 0 | Vimentin | SYVTTSTR | 914.4578 | 0.1640316 |
| 14B | 925.40149 | 1 | 0 | Vimentin | QESNEYR | 925.400996 | 0.5338226 |
| 15B | 932.4679 | 1 | 0 | Vimentin | LLEGEESR | 932.4684 | -0.536211 |
| 1C | 933.47058 | 1 | 1 | Vimentin | LLEGEESR | 933.47125 | -0.717751 |
| 2C | 936.43921 | 1 | 0 | Vimentin | SYVTTSTR+Na | 936.4396 | -0.416471 |
| 3C | 1046.5313 | 1 | 0 | Vimentin | LQEEMLQR | 1046.5299 | 1.2899775 |
| 4C | 1076.5037 | 1 | 0 | Vimentin | DNLAEDIMR | 1076.5041 | -0.40873 |
| 5C | 1081.5034 | 1 | 0 | Vimentin | QESNEYRR | 1081.5021 | 1.2205247 |
| 6C | 1082.5039 | 1 | 1 | Vimentin | QESNEYRR | 1082.50486 | -0.877594 |
| 7C | 1088.5322 | 1 | 0 | Vimentin | QDVDNASLAR | 1088.5331 | -0.799241 |
| 8C | 1089.5403 | 1 | 1 | Vimentin | QDVDNASLAR | 1089.53587 | 4.0475951 |
| 9C | 1093.5281 | 1 | 0 | Vimentin | FADLSEAANR | 1093.527276 | 0.7352354 |
| 10C | 1094.5305 | 1 | 1 | Vimentin | FADLSEAANR | 1094.53013 | 0.3563173 |
| 11C | 1095.5366 | 1 | 2 | Vimentin | FADLSEAANR | 1095.53272 | 3.5599119 |
| 12C | 1095.5449 | 1 | 0 | Vimentin | TSSPGGVYATR (semitrypti | 1095.5429 | 1.8438347 |
| 13C | 1096.5463 | 1 | 1 | Vimentin | TSSPGGVYATR (semitrypti | 1096.54578 | 0.4377382 |
| 14C | 1115.509 | 1 | 0 | Vimentin | FADLSEAANR+Na | 1115.5091 | -0.062752 |
| 15C | 1115.5687 | 1 | 0 | Vimentin | VELQELNDR | 1115.569176 | -0.399796 |
| 1D | 1116.5134 | 1 | 1 | Vimentin | FADLSEAANR+Na | 1118.527723 | -1800.843 |
| 2D | 1116.574 | 1 | 1 | Vimentin | VELQELNDR | 1116.572 | 1.7643287 |
| 3D | 1117.5772 | 1 | 2 | Vimentin | VELQELNDR | 1117.57459 | 2.2906748 |
| 4D | 1125.6047 | 1 | 0 | Vimentin | FANYIDKVR | 1125.6051 | -0.319828 |
| 5D | 1126.6099 | 1 | 1 | Vimentin | FANYIDKVR | 1126.60804 | 1.6154687 |
| 6D | 1137.5519 | 1 | 0 | Vimentin | VELQELNDR+Na | 1137.5509 | 0.8614999 |
| 7D | 1138.5549 | 1 | 1 | Vimentin | VELQELNDR+Na | 1138.553943 | 0.8668891 |
| 8D | 1245.6947 | 1 | 0 | Vimentin | SLGSALRPTTSR (semitrypt | 1245.690976 | 2.9895055 |
| 9D | 1246.6941 | 1 | 1 | Vimentin | SLGSALRPTTSR (semitrypt | 1246.69376 | 0.2647001 |
| 10D | 1254.5715 | 1 | 0 | Vimentin | LGDLYEEEMR | 1254.5671 | 3.5310985 |
| 11D | 1255.571 | 1 | 1 | Vimentin | LGDLYEEEMR | 1255.57 | 0.8283091 |

| | | | | | | | |
|---|---|---|---|---|---|---|---|
| 12D | 1270.5642 | 1 | 0 | Vimentin | LGDLYEEEMR+Oxidation | 1270.562 | 1.7393878 |
| 13D | 1271.5691 | 1 | 1 | Vimentin | LGDLYEEEMR+Oxidation | 1271.56492 | 3.2794236 |
| 14D | 1295.6685 | 1 | 0 | Vimentin | MALDIEIATYR | 1295.6664 | 1.5899154 |
| 15D | 1296.6091 | 1 | 0 | Vimentin | EEAESTLQSFR | 1296.606676 | 1.8926326 |
| 1E | 1297.6119 | 1 | 1 | Vimentin | EEAESTLQSFR | 1297.60954 | 1.8495548 |
| 2E | 1298.6162 | 1 | 2 | Vimentin | EEAESTLQSFR | 1298.61219 | 3.0956124 |
| 3E | 1309.606 | 1 | 0 | Vimentin | NLQEAEEWYK | 1309.6059 | 0.0458153 |
| 4E | 1318.5908 | 1 | 0 | Vimentin | EEAESTLQSFR+Na | 1318.5884 | 1.835296 |
| 5E | 1319.5956 | 1 | 1 | Vimentin | EEAESTLQSFR+Na | 1319.591483 | 3.1047487 |
| 6E | 1331.5872 | 1 | 0 | Vimentin | NLQEAEEWYK+Na | 1331.5877 | -0.405531 |
| 7E | 1353.7002 | 1 | 0 | Vimentin | TYSLGSALRPTTS(semitrypt | 1353.7009 | -0.517101 |
| 8E | 1354.7053 | 1 | 1 | Vimentin | TYSLGSALRPTTS(semitrypt | 1354.70378 | 1.1367799 |
| 9E | 1423.7661 | 1 | 0 | Vimentin | MALDIEIATYRK | 1423.7614 | 3.3081386 |
| 10E | 1424.7653 | 1 | 1 | Vimentin | MALDIEIATYRK | 1424.76428 | 0.6878331 |
| 11E | 1439.7573 | 1 | 0 | Vimentin | MALDIEIATYRK+oxidation | 1439.7563 | 0.7084532 |
| 12E | 1465.7763 | 1 | 0 | Spondin-1 | AQWPAWQPLNVR | 1465.769876 | 4.348568 |
| 13E | 1472.6857 | 1 | 0 | | | | |
| 14E | 1472.741 | 1 | 0 | Vimentin | TLYTSSPGGVYATR | 1472.737976 | 2.0329482 |
| 15E | 1472.7518 | 1 | 0 | Ras-related protein Ra | VNLAIWDTAGQER | 1472.749176 | 1.802072 |
| 1F | 1473.6882 | 1 | 1 | | | | |
| 2F | 1473.7448 | 1 | 1 | Vimentin | TLYTSSPGGVYATR | 1473.74092 | 2.5988286 |
| 3F | 1473.7556 | 1 | 1 | Ras-related protein Ra | VNLAIWDTAGQER | 1473.75209 | 2.3952468 |
| 4F | 1473.76 | 1 | 0 | | | | |
| 5F | 1474.75 | 1 | 2 | Vimentin | TLYTSSPGGVYATR | 1474.74365 | 4.3058331 |
| 6F | 1475.7532 | 1 | 3 | Vimentin | TLYTSSPGGVYATR | 1475.74629 | 4.662048 |
| 7F | 1494.7202 | 1 | 0 | Vimentin | TLYTSSPGGVYATR+Na | 1494.7198 | 0.2742989 |
| 8F | 1495.7273 | 1 | 1 | Vimentin | TLYTSSPGGVYATR+Na | 1495.722863 | 2.9597729 |
| 9F | 1496.7325 | 1 | 2 | Vimentin | TLYTSSPGGVYATR+Na | 1496.725593 | 4.6414654 |
| 10F | 1508.7312 | 1 | 0 | Vimentin | MFGGPGTASRPSSTR | 1508.727476 | 2.4683053 |
| 11F | 1509.7346 | 1 | 1 | Vimentin | MFGGPGTASRPSSTR | 1509.73021 | 2.9210517 |
| 12F | 1509.8064 | 1 | 0 | Vimentin | TYSLGSALRPTTSR | 1509.801976 | 2.9301856 |
| 13F | 1509.8224 | 1 | 0 | | | | |
| 14F | 1510.752 | 1 | 0 | Laminin subunit gamn | QDIAVISDSYFPR | 1510.7536 | -1.09217 |
| 15F | 1510.8101 | 1 | 1 | Vimentin | TYSLGSALRPTTSR | 1510.80483 | 3.4617311 |
| 1G | 1510.8203 | 1 | 0 | | | | |
| 2G | 1510.8272 | 1 | 1 | | | | |
| 3G | 1511.814 | 1 | 2 | Vimentin | TYSLGSALRPTTSR | 1511.8075 | 4.2730308 |
| 4G | 1512.818 | 1 | 3 | Vimentin | TYSLGSALRPTTSR | 1512.81009 | 5.2220699 |
| 5G | 1521.8081 | 1 | 0 | Spectrin beta, non-ery | HALVEADIGIQAER | 1521.802 | 4.014977 |
| 6G | 1522.813 | 1 | 1 | Spectrin beta, non-ery | HALVEADIGIQAER | 1522.80484 | 5.3322657 |
| 7G | 1524.726 | 1 | 0 | Vimentin | MFGGPGTASRPSSTR+Oxid | 1524.7224 | 2.3282927 |
| 8G | 1525.7285 | 1 | 1 | Vimentin | MFGGPGTASRPSSTR+Oxid | 1525.72512 | 2.2284486 |
| 9G | 1526.7336 | 1 | 2 | Vimentin | MFGGPGTASRPSSTR+Oxid | 1526.72669 | 4.5522228 |
| 10G | 1531.7875 | 1 | 0 | Vimentin | TYSLGSALRPTTSR+Na | 1531.7838 | 2.4024278 |
| 11G | 1532.79 | 1 | 1 | Vimentin | TYSLGSALRPTTSR+Na | 1532.786773 | 2.1314119 |
| 12G | 1533.7952 | 1 | 2 | Vimentin | TYSLGSALRPTTSR+Na | 1533.789443 | 3.7338893 |
| 13G | 1539.9183 | 1 | 0 | Vimentin | ILLAELEQLKGQGK | 1539.9105 | 5.0847111 |
| 14G | 1540.7571 | 1 | 0 | Vimentin | ETNLDSLPLVDTHS | 1540.749276 | 5.0650681 |
| 15G | 1561.8958 | 1 | 0 | Vimentin | ILLAELEQLKGQGK+Na | 1561.8923 | 2.2088591 |
| 1H | 1562.7378 | 1 | 0 | Vimentin | ETNLDSLPLVDTHS+Na | 1562.7307 | 4.5369301 |
| 2H | 1570.8992 | 1 | 0 | Vimentin | ISLPLPNFSSLNLR | 1570.895176 | 2.5424994 |
| 3H | 1571.7263 | 1 | 0 | | | | |
| 4H | 1571.9004 | 1 | 1 | Vimentin | ISLPLPNFSSLNLR | 1571.89808 | 1.4695609 |
| 5H | 1572.7312 | 1 | 1 | | | | |
| 6H | 1572.9031 | 1 | 2 | Vimentin | ISLPLPNFSSLNLR | 1572.90086 | 1.4114049 |
| 7H | 1573.9094 | 1 | 3 | Vimentin | ISLPLPNFSSLNLR | 1573.90355 | 3.7295805 |
| 8H | 1587.8042 | 1 | 0 | Vimentin | TNEKVELQELNDR | 1587.7973 | 4.3456429 |
| 9H | 1588.8065 | 1 | 1 | Vimentin | TNEKVELQELNDR | 1588.80281 | 2.2910332 |
| 10H | 1592.8828 | 1 | 0 | Vimentin | ISLPLPNFSSLNLR+Na | 1592.877 | 3.6474882 |

| | | | | | | | |
|---|---|---|---|---|---|---|---|
| 11H | 1593.8894 | 1 | 1 | Vimentin | ISLPLPNFSSLNLR+Na | 1593.880023 | 5.8831279 |
| 12H | 1637.9033 | 1 | 0 | | | | |
| 13H | 1638.9087 | 1 | 1 | | | | |
| 14H | 1659.9119 | 1 | 0 | | | | |
| 15H | 1660.9181 | 1 | 1 | | | | |
| 1I | 1665.907 | 1 | 0 | | | | |
| 2I | 1666.9108 | 1 | 1 | | | | |
| 3I | 1668.8539 | 1 | 0 | Vimentin | ETNLDSLPLVDTHSK | 1668.8439 | 5.9801878 |
| 4I | 1669.8557 | 1 | 1 | Vimentin | ETNLDSLPLVDTHSK | 1669.84683 | 5.317853 |
| 5I | 1670.856 | 1 | 2 | Vimentin | ETNLDSLPLVDTHSK | 1670.84956 | 3.8303867 |
| 6I | 1671.8574 | 1 | 3 | Vimentin | ETNLDSLPLVDTHSK | 1671.85219 | 3.128267 |
| 7I | 1690.8329 | 1 | 0 | Vimentin | ETNLDSLPLVDTHSK+Na | 1690.8257 | 4.2346174 |
| 8I | 1691.8369 | 1 | 1 | Vimentin | ETNLDSLPLVDTHSK+Na | 1691.828773 | 4.7977668 |
| 9I | 1692.8408 | 1 | 2 | Vimentin | ETNLDSLPLVDTHSK+Na | 1692.831503 | 5.4919819 |
| 10I | 1701.9128 | 1 | 0 | Tubulin alpha-1D chai | AVFVDLEPTVIDEVR | 1701.905776 | 4.1506411 |
| 11I | 1706.8046 | 1 | 0 | | | | |
| 12I | 1712.8167 | 1 | 0 | | | | |
| 13I | 1713.8235 | 1 | 1 | | | | |
| 14I | 1720.8381 | 1 | 0 | AHNAK nucleoprotein | VDIDTPDIDIHGPEGK | 1720.838776 | -0.375398 |
| 15I | 1721.8457 | 1 | 1 | AHNAK nucleoprotein | VDIDTPDIDIHGPEGK | 1721.84176 | 2.2882474 |
| 1J | 1728.7842 | 1 | 0 | | | | |
| 2J | 1729.7905 | 1 | 1 | | | | |
| 3J | 1734.8215 | 1 | 0 | Vimentin | LQDEIQNMKEEMAR | 1734.8149 | 3.8217334 |
| 4J | 1735.8263 | 1 | 1 | Vimentin | LQDEIQNMKEEMAR | 1735.81774 | 4.9141104 |
| 5J | 1896.7441 | 1 | 0 | | | | |
| 6J | 1897.752 | 1 | 1 | | | | |
| 7J | 1898.7339 | 1 | 0 | | | | |
| 8J | 2126.084 | 1 | 0 | Vimentin | LLQDSVDFSLADAINTEFK | 2126.06519 | 8.8379228 |
| 9J | 2127.0853 | 1 | 1 | Vimentin | LLQDSVDFSLADAINTEFK | 2127.06818 | 8.0627411 |
| 10J | 2128.0906 | 1 | 2 | Vimentin | LLQDSVDFSLADAINTEFK | 2128.07101 | 9.1961217 |
| 11J | 2148.0647 | 1 | 0 | Spectrin alpha, non-er | ELPTAFDYVEFTRSLFVN | 2148.0648 | -0.046554 |
| 12J | 2149.0664 | 1 | 1 | Spectrin alpha, non-er | ELPTAFDYVEFTRSLFVN | 2149.0678 | -0.646792 |
| 13J | 2150.0693 | 1 | 2 | Spectrin alpha, non-er | ELPTAFDYVEFTRSLFVN | 2150.07067 | -0.618584 |
| 14J | 2202.9706 | 1 | 0 | Vimentin | EMEENFSVEAANYQDTIGR | 2202.9608 | 4.4304011 |
| 15J | 2203.9745 | 1 | 1 | Vimentin | EMEENFSVEAANYQDTIGR | 2203.96369 | 4.8821131 |

Supplementary Table 3: The summarized result for peptide identification.

| Protein | Number of signals identified | | | |
|---|---|---|---|---|
| | OC_equatorial | IC_equatorial | OC_axial | IC_axial |
| Vimentin | 112 | | 69 | |
| Filensin | | 38 | | 54 |
| CP49 | | 52 | | 44 |
| Spectrin beta, non-erythrocytic 1 | 2 | | 6 | 2 |
| Spectrin alpha, non-erythrocytic 1 | 3 | 6 | 3 | 2 |
| Ras-related protein Rab-21 | 2 | | | |
| Laminin subunit gamma 1 | 1 | | | |
| AHNAK nucleoprotein | 2 | | | |
| Spondin-1 | 1 | | 1 | |
| Tubulin alpha-1D chain | 1 | | 1 | |
| Tubulin beta-2B/2C chain | | | 2 | |
| Actin, cytoplasmic 1/2 | | 1 | 4 | |
| Heat shock cognate 71 kDa protein | | 1 | | |
| Peptidyl-prolyl cis-trans isomerase FKBP3 | | 1 | 2 | |
| Ankynin-2 | | | 2 | |
| Ankyrin-3 | | | | 1 |
| Fructose-bisphosphate aldolase | | | 1 | |
| Guanine nucleotide-binding protein subunit γ | | | 2 | |
| Monocarboxylate transporter 1 | | | 1 | |
| Reticulon | | | 1 | 1 |
| Alpha-crystallin A chain | | | | 2 |
| Gamma-crystallin B/A | | | | 1 |
| Peroxiredoxin-1 | | | | 3 |
| Plectin | | | | 1 |
| Protein 4.1 | | | | 2 |
| Unidentified | 26 | 51 | 55 | 37 |

Supplementary Figure 1: Images of top 150 signals colocalized with the outer cortex (axial section)

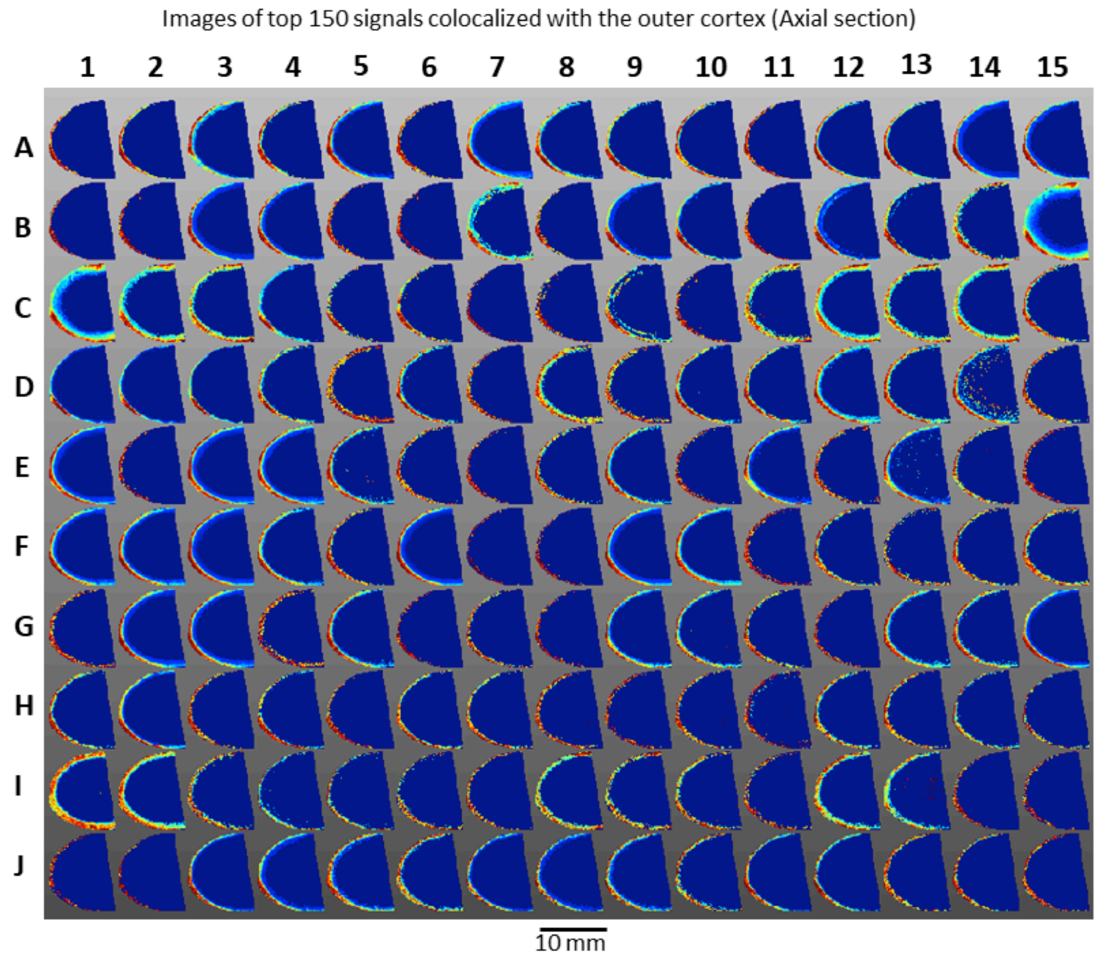

Supplementary Figure 2: Images of top 150 signals colocalized with the outer cortex (equatorial section)

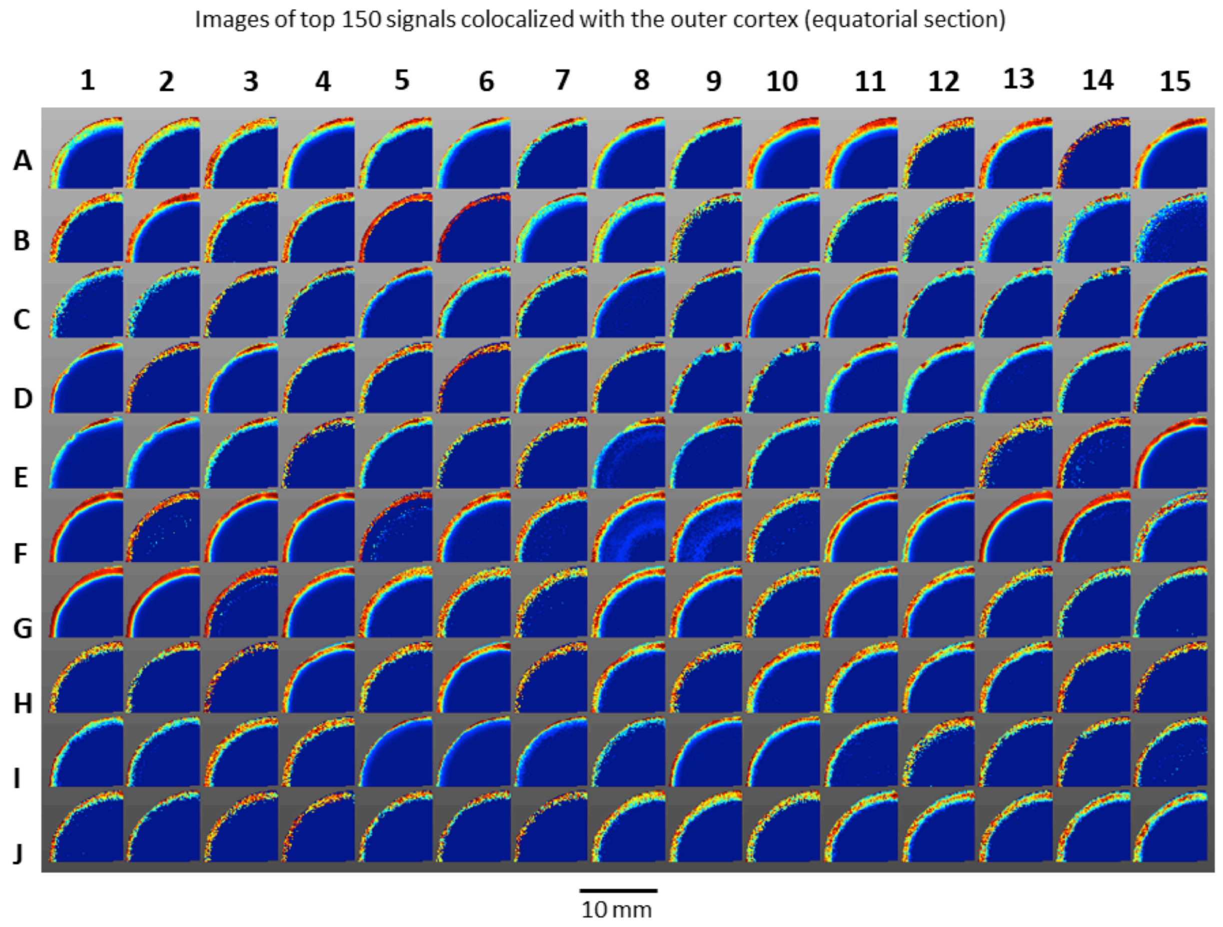

Supplementary Figure 3: Images of top 150 signals colocalized with the inner cortex (axial section)

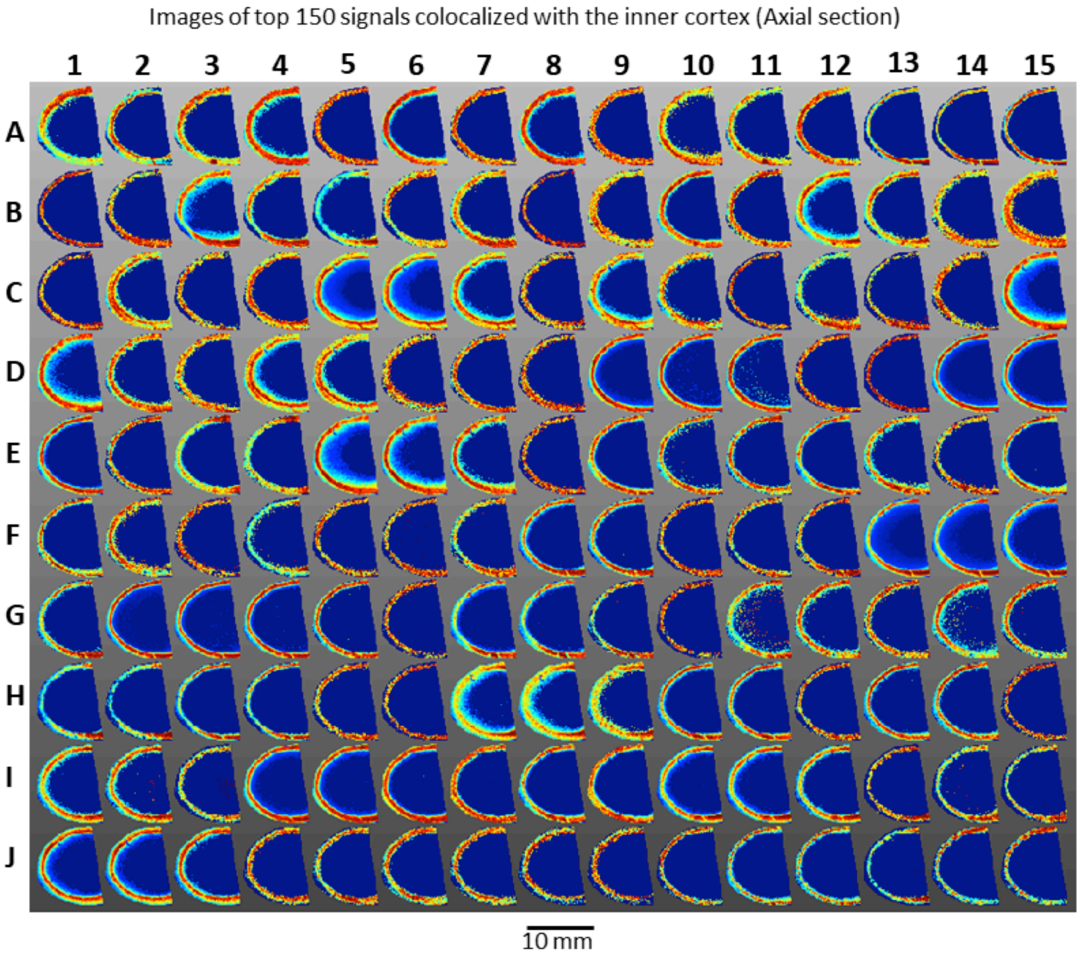

Supplementary Figure 4: Images of top 150 signals colocalized with the inner cortex (equatorial section)

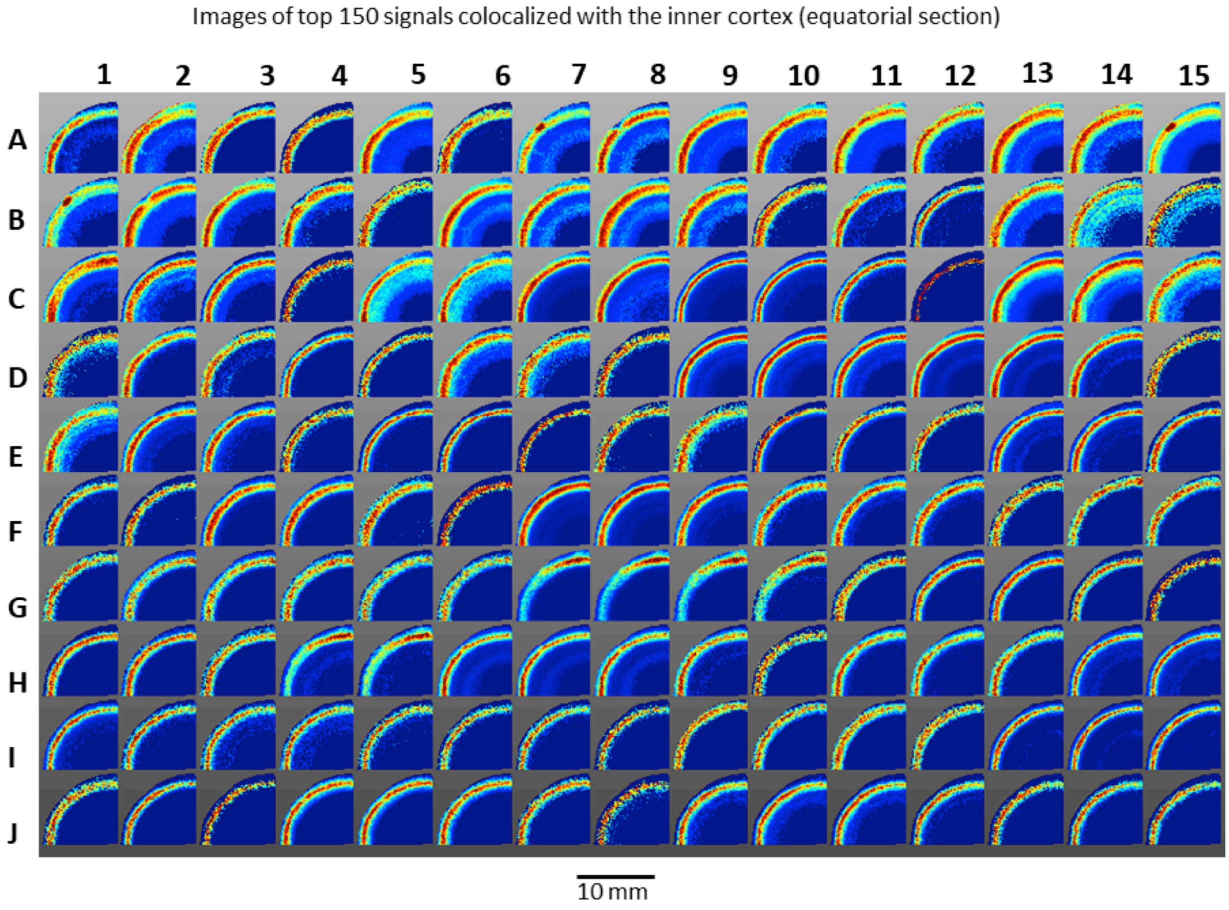

Supplementary Figure 5: Tandem mass spectrum for N-acetylated vimentin peptide 51-64.

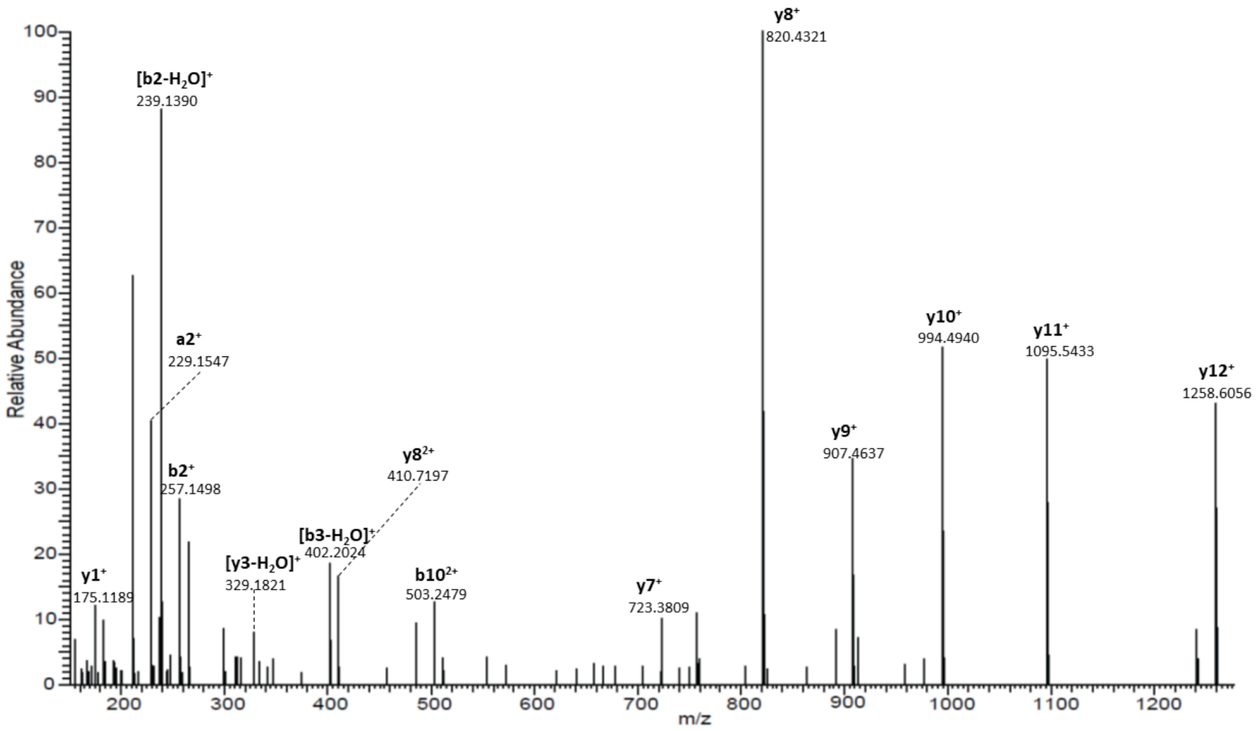
